# Learning with uncertainty for biological discovery and design

**DOI:** 10.1101/2020.08.11.247072

**Authors:** Brian Hie, Bryan Bryson, Bonnie Berger

## Abstract

Machine learning that generates biological hypotheses has transformative potential, but most learning algorithms are susceptible to pathological failure when exploring regimes beyond the training data distribution. A solution is to quantify prediction *uncertainty* so that algorithms can gracefully handle novel phenomena that confound standard methods. Here, we demonstrate the broad utility of robust uncertainty prediction in biological discovery. By leveraging Gaussian process-based uncertainty prediction on modern pretrained features, we train a model on just 72 compounds to make predictions over a 10,833-compound library, identifying and experimentally validating compounds with nanomolar affinity for diverse kinases and whole-cell growth inhibition of *Mycobacterium tuberculosis*. We show how uncertainty facilitates a tight iterative loop between computation and experimentation, improves the generative design of novel biochemical structures, and generalizes across disparate biological domains. More broadly, our work demonstrates that uncertainty should play a key role in the increasing adoption of machine learning algorithms into the experimental lifecycle.

## Introduction

As unprecedented high-throughput assays continue to transform biology [1]–[6], the ultimate goal of these studies remains the same: to generate hypotheses that elucidate important features of biological systems [7], [8]. The growing volume of experimental data underscores the importance of robust, systematic strategies to explore these results and identify experimental conditions that give rise to a desirable biological outcome.

Machine learning algorithms offer a way to translate existing data into actionable biological hypotheses [9]–[14]. However, while hypothesis generation often relies on a human expert’s intuitive certainty or uncertainty about a given hypothesis, this intuition is not automatically built into a machine learning algorithm, making these algorithms susceptible to overconfident predictions, especially when access to training data is limited [15], [16]. Instead, an intelligent algorithm that quantifies prediction uncertainty [17]–[21] could help focus experimental effort on hypotheses with a high likelihood of success, which is especially useful when new data acquisition is slow or arduous; or, the algorithm could alert a researcher to experiments with greater novelty but also with a greater risk of failure [17].

While uncertainty is gradually becoming recognized as a critical property in learning algorithms [22]–[24], in the biological setting, most machine learning studies simply do not consider uncertainty, while those that do are limited to specific tasks or *in silico* validation [25]–[29]. Given the growing interest in applying machine learning to biology, here we comprehensively demonstrate the benefit of learning with uncertainty and highlight a general, practical way to do so. One of our key methodological findings is that a class of algorithms based on Gaussian processes (GPs) [18]–[21], trained on rich and modern features, provides useful quantification of uncertainty while also enabling substantive biological discoveries even with a limited amount of training data.

Our main discovery task involves selecting small molecules based on predicted binding affinity with mycobacterial and human kinases, a fundamental biochemical and pharmacological task. Leveraging the robustness offered by prediction uncertainty, we train our models with information from just 72 kinase inhibitors [1], perform an *in silico* screen of an unbiased 10,833-compound chemical library [30], and, crucially, experimentally validate our machine-generated hypotheses with *in vitro* binding assays. Remarkably, the GP-based models with a principled consideration of uncertainty acquire a set of interactions that is unprecedented for compound-target interaction prediction, with a hit rate of 90% and with highly potent affinities in the nanomolar or sub-nanomolar range involving all of our tested kinases, including novel interactions that are uncharacterized by previous studies.

A key benefit of our approach is that it facilitates a more interactive experience between the human researcher and the learning algorithm. In particular, uncertainty prediction helps guide iterative rounds of computation and experimentation that progressively explore unknown biological search spaces [28], [31]. To illustrate this concept, we make uncertainty-based predictions, experimentally validate those predictions, incorporate the new data into a second prediction round, and discover another compound with potent nanomolar affinity for PknB, an essential *Mycobacterium tuberculosis* (Mtb) kinase, with low Tanimoto similarity to any compound in the training set.

We also demonstrate how robust, machine-generated hypotheses can spawn broader lines of research by showing that a subset of our high affinity compounds leads to whole cell growth inhibition of Mtb, the leading cause of infectious disease death globally [32]. We further use uncertainty to improve the generative design of novel small molecule structures based on high biochemical affinity for a given target. Our work demonstrates how uncertainty-based prediction can aid and augment the cycle of hypothesis generation and data collection at the heart of the scientific method.

To illustrate the generality of our approach, we apply our same framework to a different task relevant to protein engineering. We show that uncertainty can improve models of the fitness landscape of *Aequorea victoria* green fluorescent protein (avGFP) [2], a workhorse tool in biology. Using just a small amount of training data, we use uncertainty to prioritize combinatorial mutants to avGFP based on predicted fluorescence, revealing important structural elements underlying preserved or even enhanced fluorescence.

We find that principled prior uncertainty, which we implement with sample-efficient GPs trained on modern features, improves learning-based prediction across a breadth of biological tasks and domains. Our results demonstrate that uncertainty should be a critical consideration as biological studies increasingly blend computational and experimental methods. This paper provides a roadmap for how researchers might better incorporate learning algorithms into the experimental life cycle.

## Results

### Uncertainty prediction enables robust machine-guided discovery: Theory and a conceptual use case

The critical concept in our study is that prediction “uncertainty” can help machine learning algorithms more robustly explore novel biological hypotheses (**Fig. 1A**) [17]–[21]. For example, consider the setting where a researcher is interested in finding a small molecule that inhibits a kinase, a problem of biochemical and pharmacological importance. When a researcher considers new inhibitors, some chemical structures might be similar to well-studied structures, and therefore might also have similar behavior. However, there is an enormous space of chemical structures with uncertain or unknown biochemistry. While notions of biochemical “similarity” or “uncertainty” might be obvious to a human expert, a standard machine learning algorithm has no corresponding notion of uncertainty, potentially leading to biased, overconfident, or pathological predictions [15], [16], [22], [23], [33]–[35].

**Figure 1:**
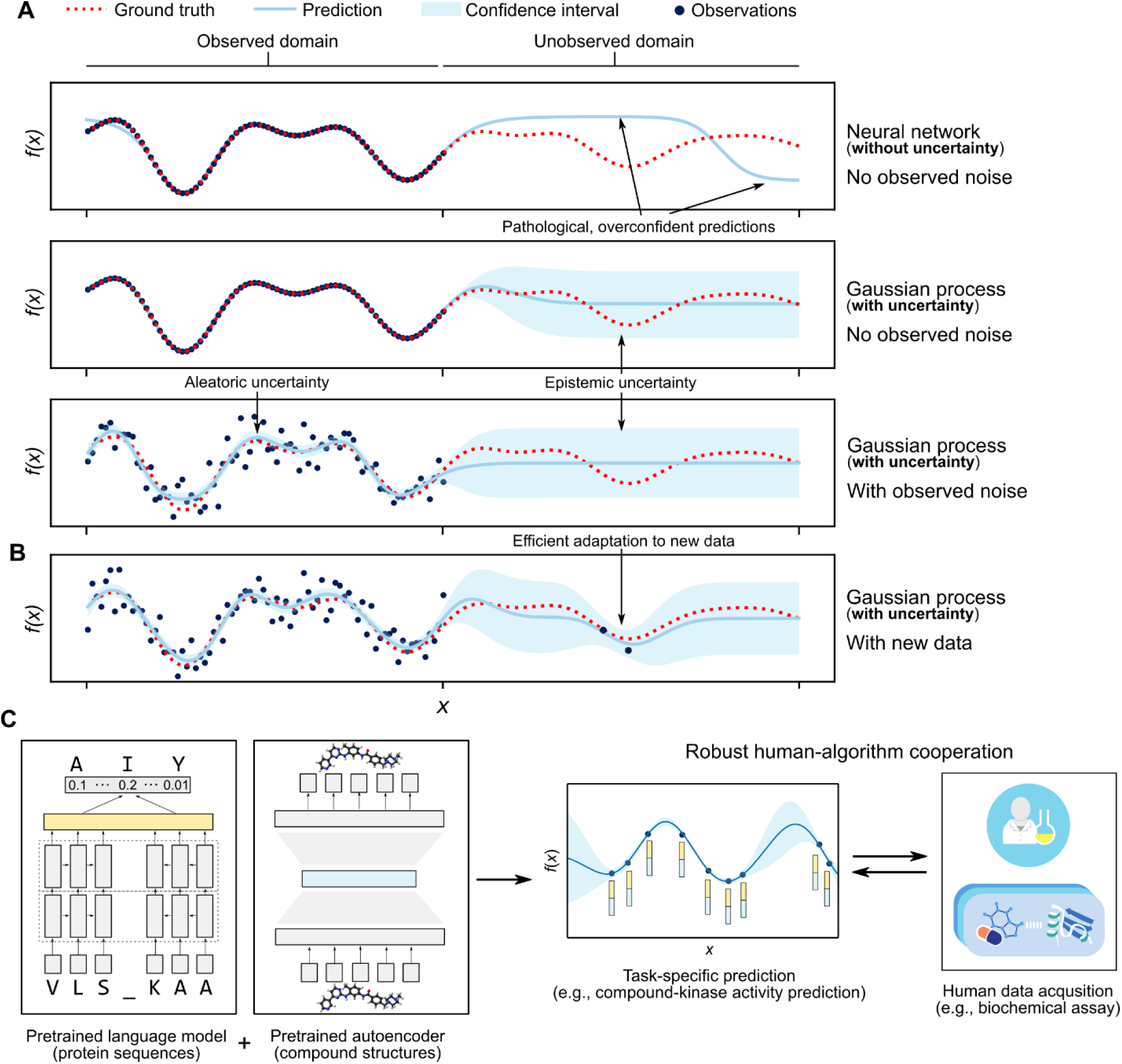
Robust uncertainty prediction for machine-guided discovery. (**A**) When a machine learning model encounters an example like nothing in its training set, its behavior is usually undefined. A way to improve robustness is for the model to report high uncertainty on such examples. Rather than output a single point prediction for each example in a given domain, more robust methods, such as a Gaussian process (GP), model the aleatoric (or statistical) uncertainty of observations and the epistemic (or systematic) uncertainty that comes from a lack of data. In a GP, the epistemic uncertainty of unexplored regions of the domain is explicitly encoded as a prior probability. (**B**) GPs can readily update their beliefs with just a handful of new datapoints. (**C**) Using modern, neural pretrained feature representations, a GP can achieve state-of-the-art prediction performance even with limited data. Knowing uncertainty helps guide a researcher when prioritizing experiments and, when combined with sample efficiency, enables a tight feedback loop between human data acquisition and algorithmic prediction.

Two additional concepts also help to improve the practicality and performance of uncertainty prediction for biological discovery. The first, “sample efficiency” (**Fig. 1B**) [19]–[21], is the ability to make use of and quickly adapt to new data. Sample efficiency is especially critical in domains where new data collection is limited or slow (for example, synthesizing and testing customized small-molecule drugs [36]). The second concept is the notion of “pretraining” [11], [37]. Pretraining automatically extracts meaningful, general features in a task-agnostic, or an unsupervised, way (for example, pretrained features could be extracted from a large database of chemical structures or protein sequences without information on compound-protein interactions). Pretrained features can subsequently improve the performance of uncertainty prediction within a more specific downstream task (**Fig. 1C**).

GPs are prime candidates for machine learning-based hypothesis generation since they naturally quantify prediction uncertainty [18], are highly sample-efficient [21], and can readily incorporate pretrained features. GPs allow a researcher to specify high uncertainty when the training distribution provides little information on unseen test examples, which is referred to as *epistemic* uncertainty (**Fig. 1**) [17]. For example, when predicting compound-kinase affinity, a reasonable prior uncertainty would assign most of the probability to low affinity but still assign a small probability to high affinity. Rather than outputting a single point prediction for each datapoint, GPs output a *probability distribution* (i.e., a Gaussian distribution), where a location-related statistic, like the mean, can be used as the prediction value and a dispersion-related statistic, like the standard deviation, can be used as the uncertainty score.

As datapoints become more distal to the training set, GP uncertainty also grows to approach the prior uncertainty, analogous to human uncertainty increasing on examples that deviate from existing knowledge (**Fig. 1**) [17]–[21]. A researcher can then use an *acquisition function* to select predictions with both good predictive scores and low uncertainty for further experimental validation. A straightforward and widely-used acquisition function, called the upper confidence bound (UCB), adds the prediction and uncertainty scores with a weight factor *β* controlling the importance of the uncertainty [17], [38]. An acquisition function with a high *β* prioritizes low uncertainty; in contrast, a low *β* deprioritizes uncertainty, and *β* = 0 ignores uncertainty (**Methods**).

### Uncertainty prediction enables robust machine-guided discovery: Application to compound-kinase affinity prediction

We therefore sought to assess, first, whether modelling uncertainty is competitive against standard baselines (i.e., we suffer no performance loss when incorporating uncertainty prediction) and, second, whether uncertainty prediction offers any real advantage when prioritizing biological hypotheses for a researcher to perform. As a test case for machine-guided discovery, we decided to initially focus on predicting binding affinities between small molecule compounds and protein kinases. We select this particular application since kinases have diverse pharmacological implications that include cancer and infectious disease therapeutics [39]–[42] and good quality compound-kinase affinity training data exists for a limited number of compounds [1].

Before committing resources to experimental validation of compound-kinase affinity predictions, we first set up an *in silico* simulation of the prediction and discovery process. We obtained a publicly-available dataset [1] containing binding affinity measurements, within a 0.1 to 10,000 nanomolar (nM) range, of the complete set of kinase-compound pairs among 72 compounds and 442 unique kinase proteins (the dataset contained 379 unique kinase genes with multiple mutational variants for some of the genes). We set up a cross-validation-based simulation by separating the known data into training and test data (**Supplementary Fig. 1A**). Importantly, to simulate out-of-distribution prediction, we ensured that approximately one-third of the test data contained interactions involving compounds not in the training data, one-third contained interactions involving kinase genes not in the training data, and one-third contained interactions involving compounds *and* kinase genes not in the training data (**Supplementary Fig. 1A**).

Our main set of benchmarking methods leverages unsupervised pretraining via state-of-the-art neural graph convolutional-based compound features (pretrained by the original study authors on ∼250K small molecule structures) [43] and neural language model-based protein sequence features (pretrained by the original study authors on ∼21M protein sequences) [44] (**Methods**). Subsequent regression models use a concatenation of these features to predict Kd binding affinities (**Fig. 2A**).

**Figure 2:**
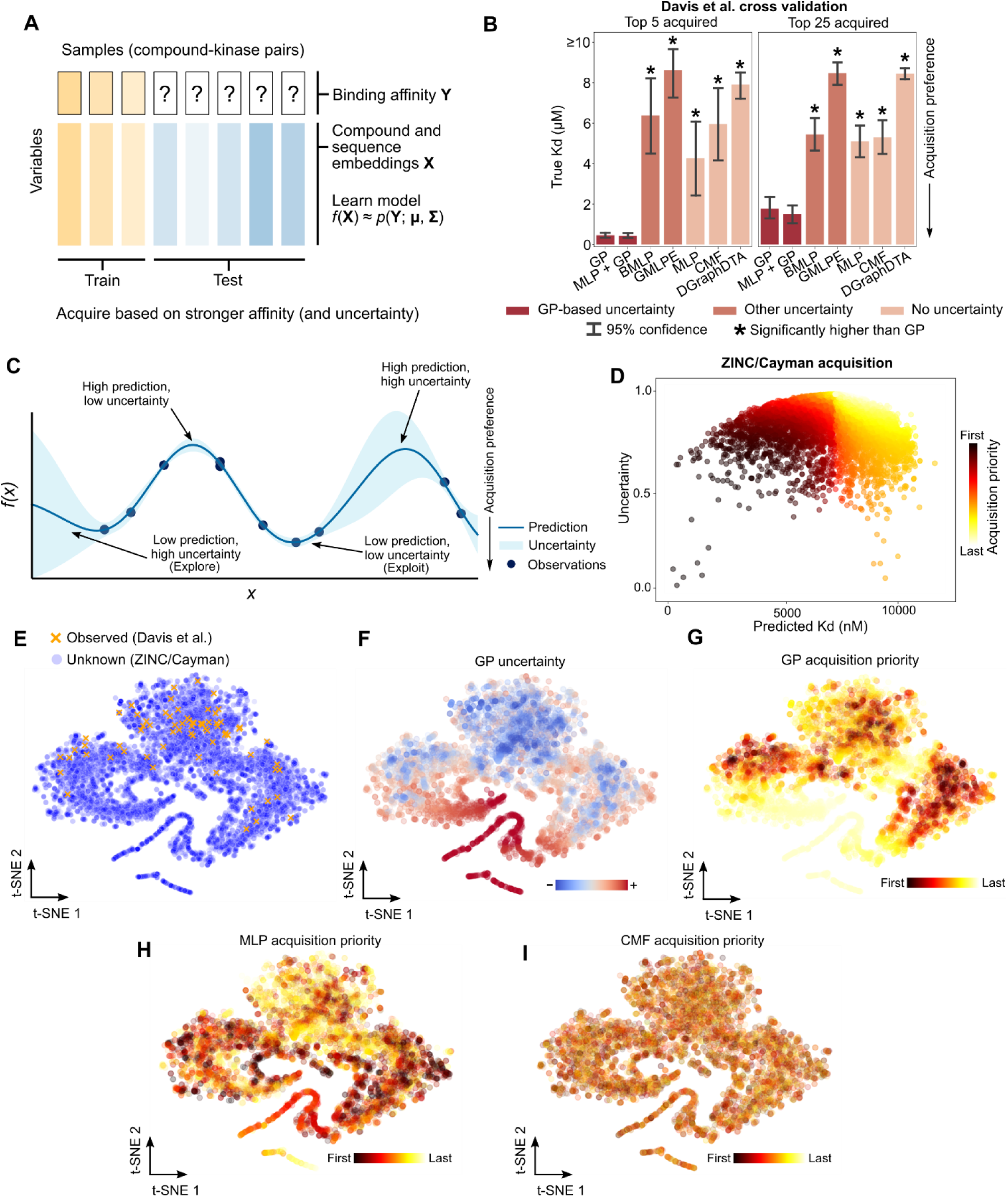
Computational prediction of compound-kinase affinity. (**A**) We desire to predict compound-kinase affinity based on features derived from compound structure and kinase sequence and use these predictions to acquire new interactions. Incorporating uncertainty into predictions is especially useful when the data distributions of the training and test sets are not guaranteed to be the same. (**B**) True Kds of the top five and twenty-five prioritized compound-kinase pairs (**Methods**) for each model over five model initialization random seeds. Bar height indicates mean Kd; statistical significance was assessed with a one-sided Welch’s *t*-test *P*-value at FDR < 0.05. (**C**) Predictions augmented with uncertainty scores enable a researcher to perform experiments in high confidence, high desirability regions (“exploitation”) or to probe potentially high desirability regions with less model confidence (“exploration”). (**D**) Each point represents a compound in the ZINC/Cayman library with an associated predicted Kd (with PknB) and uncertainty score outputted by a GP (normalized by the prior uncertainty), colored by the order the compound appears according to our acquisition function. We use an acquisition function (**Methods**) that prioritizes high confidence, low Kd predictions. (**E**) A t-SNE visualization of the compound feature space reveals regions of the compound landscape without any representative compounds with known PknB affinity measurement. (**F**) A GP assigns lower uncertainty to regions of the compound landscape close to the observed data. (**G**) A subset of the low uncertainty compounds is prioritized for experimental acquisition based on predicted binding affinity to PknB. (**H**) The MLP assigns high predicted PknB binding to a large number of out-of-distribution compounds. (**I**) CMF predictions for PknB appear to lack any meaningful structure with regards to the compound landscape. Example acquisition for other kinases is provided in **Supplementary** Fig. 3.

We benchmark methods that also learn some notion of prediction uncertainty. Our first uncertainty model fits a GP regressor [18], [45] to the training set. For a given compound-kinase pair, the GP provides a Kd prediction in the form of a Gaussian distribution, where we use the mean of the Gaussian as the prediction value and the standard deviation as the measure of uncertainty. Our second method first fits a multilayer perceptron (MLP), followed by fitting a GP to the residuals of the MLP predictions (MLP + GP) [46]. This results in a hybrid model where the prediction is the sum of the MLP and the GP estimates, while the GP variances can be used as the uncertainty scores. We also benchmark two other methods that attempt to augment neural networks with uncertainty: a Bayesian multilayer perceptron (BMLP) [23], [47] and an ensemble of MLPs that each emits a Gaussian distribution (GMLPE) [22].

We also benchmark three baseline methods without uncertainty. On the same neural pretrained features, we test an MLP, also known as a densely-connected neural network [48], [49] and collective matrix factorization (CMF), a model often used to recommend shopping items for potential buyers that can also recommend drugs for potential targets [50]–[53]; variants of both models have seen extensive use in previous compound-target interaction prediction studies. Lastly, to assess the benefit of our unsupervised pretraining-based features, we also train DGraphDTA, a graph convolutional neural network designed specifically for compound-target interaction prediction [54], from end-to-end on a simpler set of features.

The results of our cross-validation experiment using standard, average-case performance metrics show that GP-based models are consistently competitive with, and often better than, other methods based on average-case performance metrics. The Pearson correlations between the predicted Kds and the ground truth Kds for our GP and MLP + GP models over all test data are 0.35 and 0.38, respectively (*n* = 24,048 compound-kinase pairs), in contrast with 0.26, 0.23, and 0.21 for the MLP, CMF, and DGraphDTA baselines, respectively (**Supplementary Fig. 1B**). Good regression performance of GP-based methods is also consistent across all our metrics (Pearson correlation, Spearman correlation, and mean square error) when partitioning the test set based on exclusion of observed compounds, kinases, or both (**Supplementary Fig. 1B**).

Importantly, we also observed that, in this relatively data-limited training setting, rich pretrained features combined with a relatively lightweight regressor (e.g., a GP or MLP) outperformed a more complex regressor architecture (i.e., DGraphDTA) trained end-to-end on simpler features (**Supplementary Fig. 1B**). This provides evidence that pretraining with state-of-the-art unsupervised models contributes valuable information in a data-limited setting.

Where robust GP-based prediction has a substantially large advantage is in prioritizing compound-kinase pairs for further study. Importantly, in contrast to average-case metrics, focusing on top predictions directly mimics biological discovery, since researchers typically choose only a few lead predictions for further experimentation rather than testing the full, unexplored space. In GP-based models, we observed that predictions with lower uncertainty are more likely to be correct, whereas high-uncertainty predictions have worse quality (**Supplementary Fig. 1C**), allowing us to prioritize compound-kinase pairs with high predicted affinity *and* low prediction uncertainty (**Methods**). In contrast, models without uncertainty like the MLP do not distinguish confident and uncertain predictions (**Supplementary Fig. 1C**). The top compound-kinase pairs acquired by the GP-based models have strong, ground-truth affinities, while the other methods with poorly calibrated or nonexistent uncertainty quantification struggle to prioritize true interactions and acquire interactions with significantly higher Kds (**Fig. 2B** and **Supplementary Fig. 2A**). Performance of the GP-based models decreases when ignoring uncertainty (**Supplementary Fig. 2B**), suggesting that GP uncertainty helps reduce false-positives among top-acquired samples; however, other methods (BLMP and GMLPE) seem to have trouble learning meaningful uncertainty estimates (**Supplementary Fig. 2B**).

### Prioritization of compound-kinase interactions based on predicted affinity

Based on the success of our cross-validation experiments, particularly when using theoretically principled GP-based models, we then sought to perform machine learning-guided biological discovery of previously unknown compound-kinase interactions. For this task, we use all information across the pairs of 72 compounds and 442 kinases [1] as the model training data. For the search space of compounds outside of the training set, we use a collection of 10,833 compounds from the ZINC database [30] that is commercially available through the Cayman Chemical Company, allowing us to purchase high-quality compound samples for further validation experiments. Chemicals were selected solely on the basis of commercial availability, regardless of potential associations with kinases or any other biochemical property. The resulting “ZINC/Cayman library” consists of heterogeneous compounds (molecular weights range from 61 to 995 Da) with a median Morgan fingerprint Tanimoto similarity of 0.09; additional statistics for this library can be found in **Supplementary Table 1**.

In theory, out-of-distribution predictions with low predicted Kd *and* low uncertainty (**Fig. 2C,D**) will also have low *in vitro* Kds. Before validating our predictions, we first wanted to ensure that the empirical behavior of our uncertainty models matched our intuition: namely, that unknown compounds very different from any compound in the training set would also have high associated uncertainty. To do so, we visualized the 72 compounds from the training set [1] and the 10,833 unknown-affinity compounds using a two-dimensional t-distributed Stochastic Neighbor Embedding (t-SNE) [55] of the structure-based compound feature space. The embedding shows large regions of the compound landscape that are far from any compounds with known affinities (**Fig. 2E**).

We can then use a GP, trained on just 72 compounds, to assign a predicted Kd affinity (for a given kinase target) to each compound in the ZINC/Cayman library, as well as a corresponding uncertainty score. When we color the points in the visualization by uncertainty, consistent with our intuition, GP uncertainties are lower in regions near compounds with known affinities and higher in regions with many unknown-affinity compounds (**Fig. 2F**), with high correlation between the uncertainty score and test compound distance to its Euclidean nearest neighbor in the training set (Spearman *r* = 0.87, *n* = 10,833 compounds; **Methods**). The GP prioritizes compounds within the low uncertainty regimes that also have high predicted binding affinity (**Fig. 2G** and **Supplementary Fig. 3**). In contrast, the MLP assigns high priority to many compounds far from the known training examples (**Fig. 2H** and **Supplementary Fig. 3**), which is most likely due to pathological behavior on out-of-distribution examples. For comparison, CMF seems unable to learn generalizable patterns from the small number of training compounds (**Fig. 2I** and **Supplementary Fig. 3**).

### Uncertainty prediction discovers sub-nanomolar compound-kinase biochemical activity

We then performed machine-guided discovery of compound-kinase interactions. Since our *in vitro* binding assays are optimized to screen many compounds for a given kinase, we focused our validation efforts on a set of four diverse kinases: human IRAK4, a serine/threonine kinase involved in Toll-like receptor signaling [56]; human c-SRC, a tyrosine kinase and canonical proto-oncogene [57]; human p110δ, a lipid kinase and leukocytic immune regulator [58]; and Mtb PknB, a serine/threonine kinase essential to mycobacterial viability [59]. These kinases have well-documented roles in cancer, immunological, or infectious disease [39]–[42].

To compare prediction with and without uncertainty, we used either our GP or MLP models to acquire compounds from the ZINC/Cayman library with high predicted affinity for each of the four kinases of interest. We validated the top five predictions returned by the GP or MLP for each kinase using an *in vitro* biochemical assay to determine the Kd (**Methods**).

We observed that none of the predictions acquired by the MLP had a Kd of less than the top tested concentration of 10 μM (**Fig. 3** and **Supplementary Table 2**), consistent with out-of-distribution prediction resulting in pathological model bias (**Supplementary Fig. 2**). In contrast, the GP yielded 18 compound-kinase pairs with Kds less than 10 μM (out of 20 pairs tested, or a hit rate of 90%), 10 of which are lower than 100 nM (**Fig. 3** and **Supplementary Table 2**). Notably, GP acquisition yielded sub-nanomolar affinities between K252a and IRAK4 (Kd = 0.85 nM) and between PI-3065 and p110δ (Kd = 0.36 nM), automating discoveries that previously had been made with massive-scale screens or expert biochemical reasoning [42], [60]. Some compounds had predicted and validated affinities for multiple kinases, such as K252a, an member of the indolocarbazole class of compounds, many of which have broad-spectrum kinase inhibition [1]. Other compounds were only acquired for one of the kinases, including PI-3065 for p110δ, WS3 for c-SRC (Kd = 4 nM), and SU11652 for PknB (Kd = 76 nM). Interestingly, the latter two of these interactions do not seem to have existing experimental support; WS3 was developed as an inducer of pancreatic beta cell proliferation [61] and SU11652 was developed for human receptor tyrosine kinase inhibition [62].

**Figure 3:**
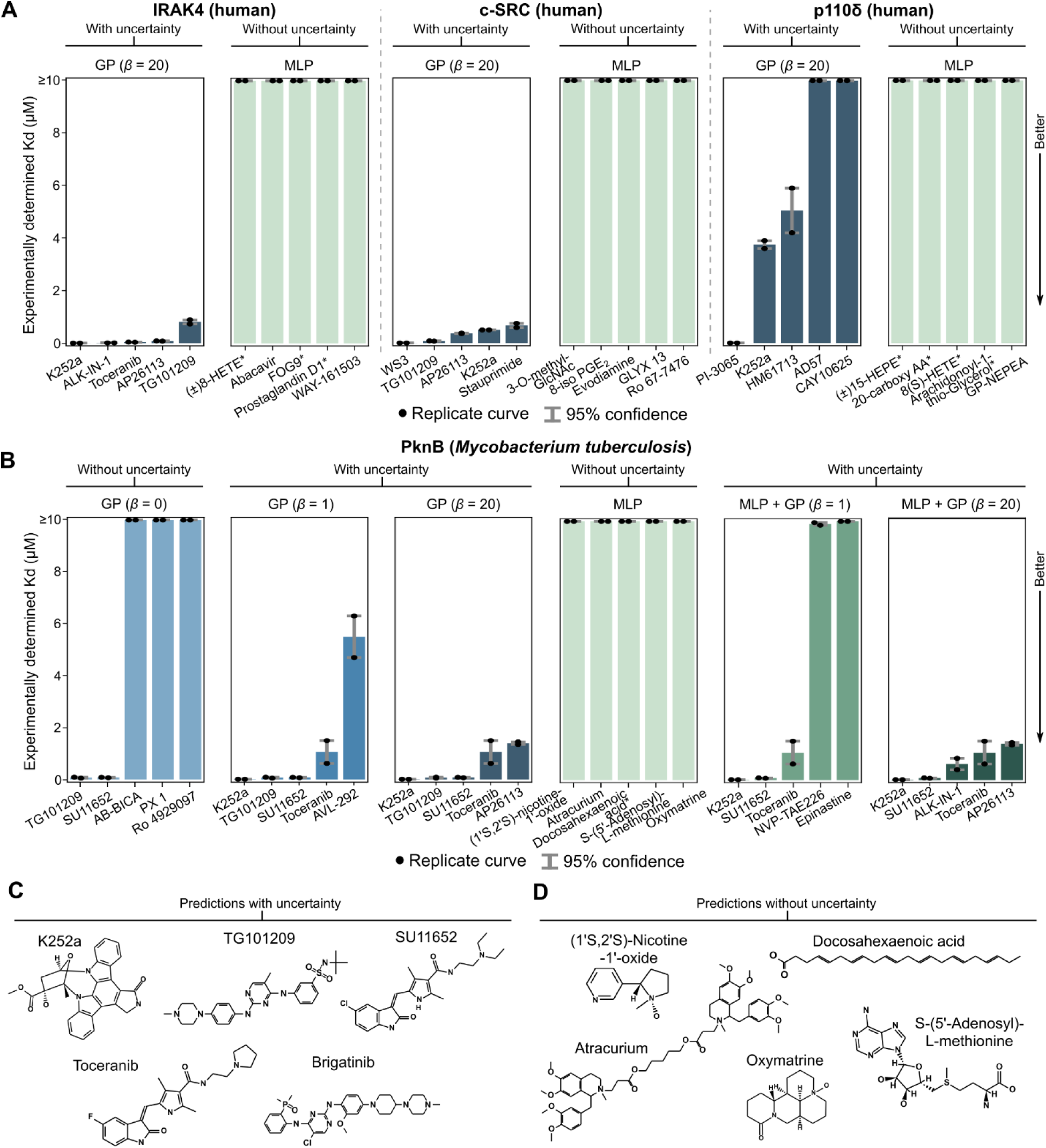
Uncertainty enables acquisition of potent compound-kinase interactions. (**A**) Binding affinity Kd for top five acquired compounds for three human kinases using a model with uncertainty (GP) and without (MLP). Asterisks after compound names indicate compounds incompatible with the validation assay (**Methods**). Mean Kd values are provided in **Supplementary Table 2**. (**B**) We validated the top five compound predictions at different acquisition *β* parameters for the models with uncertainty (GP and MLP + GP) and the top five compound predictions provided by the MLP. Incorporating uncertainty information reduces false-positive predictions. Asterisks after compound names indicate compounds incompatible with the validation assay. Mean Kd values are provided in **Supplementary Table 2**. (**C**) The structures of the compounds prioritized by the GP for PknB binding affinity with acquisition *β* = 20. (**D**) The structures of the compounds prioritized by the MLP for PknB binding affinity, none of which have strong affinity (Kd ≥ 10,000 nM).

To further assess the impact of uncertainty on prediction quality, we also performed PknB acquisition with another GP-based model (MLP + GP) and varied the weight *β* on the uncertainty (**Methods**). We validated the top five predictions from the GP and MLP + GP at *β* = 1 (tolerates some uncertainty) and *β* = 20 (prefers lowest Kds with a very low tolerance for uncertainty), as well as the top five predictions from the GP at *β* = 0 (i.e., ignoring uncertainty). At *β* = 20, the MLP + GP acquired a similarly potent set of compounds as the GP. Tolerating greater amounts of uncertainty, or ignoring it completely, led to more false-positive predictions (**Fig. 3B**).

The ability for GP-based models to yield (sub-)nanomolar affinity compound-kinase pairs in a highly out-of-distribution prediction task provides strong experimental support for our uncertainty-based learning approach. We note that training our models on information from 72 compounds to make predictions over a 10,833-compound library is a much more challenging task than any previously reported drug-target interaction prediction setting [48]–[52]. Moreover, among previous studies that performed experimental validation of machine learning-predicted compound-target interactions, none report Kds or IC/EC50s below the micromolar range [48], [52], suggesting that our approach yields most potent interactions discovered by compound-target interaction prediction to date.

A notable advantage of GP-based uncertainty quantification is that it enables an absolute assessment of prediction quality. For example, all predictions with a mean less than 10 μM (our top-tested concentration) and an interquartile range less than 2 μM resulted in true positive hits (**Supplementary Fig. 5**). In contrast, more dispersed prediction distributions had higher variability in the potency of the true binding interaction including false positives (**Supplementary Fig. 5**), suggesting that our GP-based models make better predictions when they are more confident. Uncertainty adds an interpretable dimension to machine-generated predictions; for example, a researcher with a low tolerance for false positives might ignore a generated hypothesis with a low predicted Kd but a high prediction uncertainty.

### Anti-Mtb activity of compounds with validated PknB biochemical activity

Machine-generated hypotheses also add value by stimulating follow-up biological research that provides insights beyond the initial prediction problem. For example, given the novel, potent interactions discovered by our models, we sought to further assess if the compounds had any broader relevance beyond biochemical affinity with the protein molecule itself.

PknB is a kinase that is essential to Mtb viability [59]. Bacterial kinases are less well studied than human (or other mammalian) kinases [63], [64] but are nonetheless important therapeutic targets [39], [59], [65], especially in antibiotic-resistant bacterial strains. Most importantly, tuberculosis remains the leading cause of infectious disease death globally [32], underscoring the importance of novel antibacterial therapies. Given the essentiality of PknB and our *in silico* identification of PknB-binding compounds, we sought to examine if the compounds with high binding affinity to PknB would have any impact on mycobacterial growth. This would not be guaranteed since factors like cell wall permeability or intracellular stability were not explicitly encoded in the training data.

We focused on the compounds with a Kd less than 100 nM: K252a (Kd = 11 nM), TG101209 (Kd = 71 nM), and SU11652 (Kd = 76 nM). Using the colorimetric, resazurin microtiter assay (alamar blue) [39], [66], we determined the minimum inhibitory concentration (MIC) of these compounds as well as rifampicin, a frontline antibiotic for tuberculosis [32] (**Methods**); the MICs for these compounds with H37Rv are shown in **Supplementary Table 3**. We observed that K252a and SU11652 inhibited the growth of H37Rv compared to a dimethyl sulfoxide (DMSO) vehicle control (one-sided *t*-test *P*-value of 7.0 × 10^-8^ for K252a and 3.9 × 10^-8^ for SU11652, *n =* 3 replicate cultures per condition) (**Fig. 4A** and **Supplementary Fig. 4A**).

**Figure 4:**
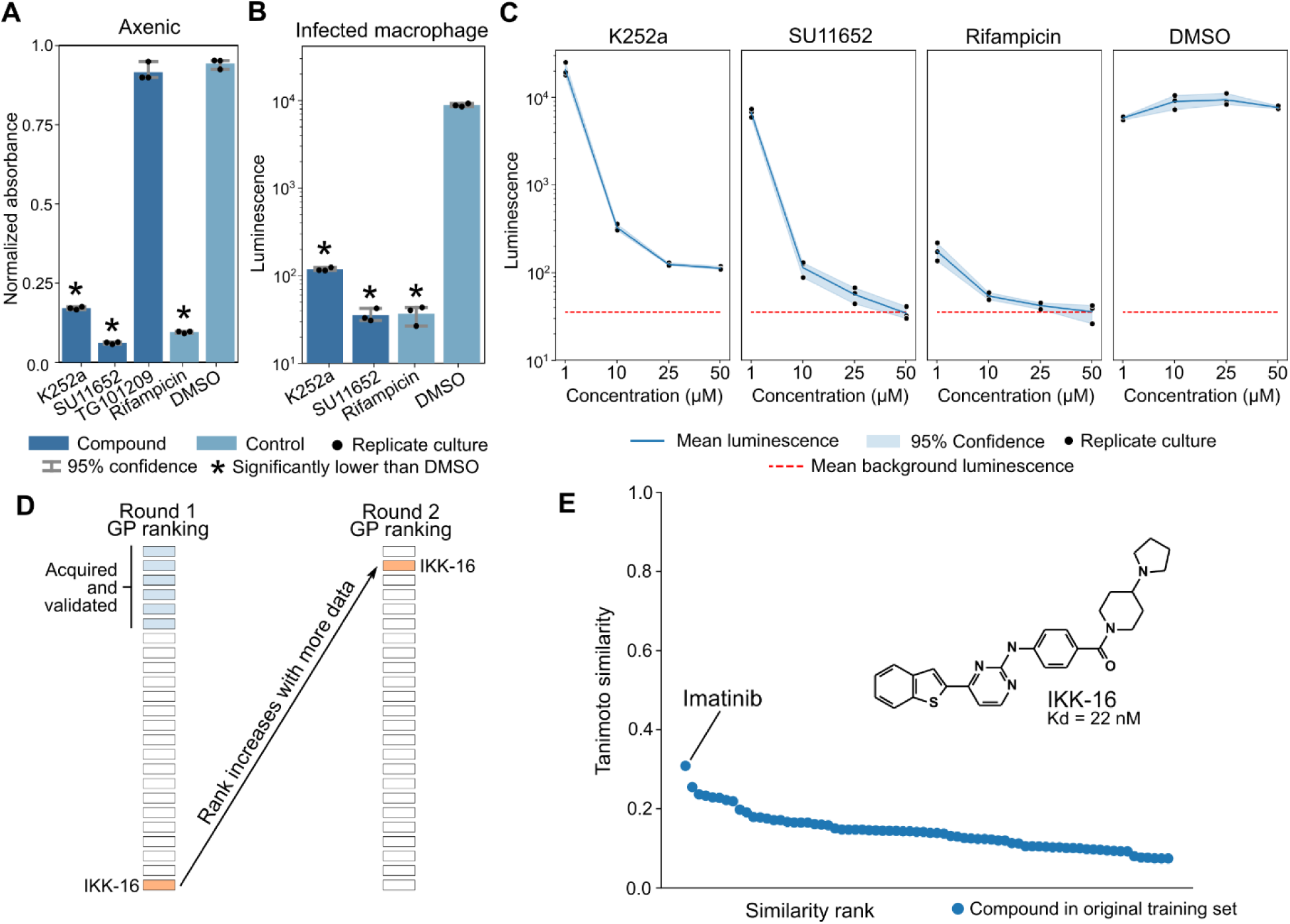
Follow-up PknB experiments reveal anti-Mtb whole cell activity and an out-of-distribution inhibitor. (**A**) Growth of axenic Mtb measured via alamar blue absorbance after five days of axenic incubation in media treated with compounds, or a DMSO vehicle control, at 50 μM. Statistical significance was assessed with a one-sided *t*-test *P*-value at FDR < 0.05. (**B**) Luminescence of luciferase-expressing Mtb from within infected human macrophages cultured in media treated with compounds at 50 μM. Statistical significance was assessed with a one-sided *t*-test *P*-value at FDR < 0.05. (**C**) Dose-response of K252a, SU11652, rifampicin, or a DMSO vehicle control on luminescence of luciferase-expressing Mtb from within infected human macrophages after five days of culture post-infection. (**D**) IKK-16 was ranked 24 by the GP during the first round of compound acquisition. Six of the compounds above IKK-16 in the first-round GP ranking were acquired for experimental validation (the sixth-ranked compound was in the top five for the MLP + GP). Following model retraining on first-round PknB binding acquisitions across all models, IKK-16 was the second-ranked compound. (**E**) All 72 compounds in the original training set have a Morgan fingerprint (radius 2, 2048 bits) Tanimoto similarity of 0.31 or less with IKK-16 (structure shown).

SU11652 is a well-documented inhibitor of human receptor tyrosine kinases including PDGFR, VEGFR, and Kit [62]. TG101209 did not inhibit growth of H37Rv (one-sided *t*-test *P*-value of 0.11, *n =* 3 replicate cultures per condition) (**Fig. 4A** and **Supplementary Fig. 4A**), perhaps due to low cell permeability [67], [68]. These results were corroborated using additional validation where Mtb expressing the *luxABCDE* cassette (luxMtb) was incubated with increasing concentrations of K252a, SU11652, and TG101209 (**Supplementary Fig. 4B**; **Methods**).

We further validated these results in a more complex, host-pathogen model. We utilized a previously described assay for monitoring intracellular growth where macrophages are infected with luxMtb and luminescence is measured as a proxy of bacterial growth [69], [70] (**Methods**). We infected macrophages with luxMtb for 4 hours prior to the addition of compounds dissolved in cell culture media. Consistent with our axenic culture experiments, treatment with K252a and SU11652 resulted in less luminescence as compared to DMSO (one-sided *t*-test *P*-value of 2.9 × 10^-6^ for K252a and 2.8 × 10^-6^ for SU11652; *n =* 3 replicate cultures per condition) (**Fig. 4B,C**). In examining the literature for prior work on compounds targeting PknB, we identified support for K252a as an inhibitor of PknB kinase activity and Mtb growth [59], [65]. These previous studies and our results nominate future experiments to further investigate the biochemistry of PknB and the potential use of K252a and SU11652 as scaffolds for PknB-related drug development. These results also illustrate how learning algorithms with uncertainty can stimulate productive follow-up hypotheses, in this case highlighting molecular structures that should be further investigated for therapeutic relevance to the leading cause of infectious disease death among humans.

### Active learning with uncertainty reveals a structurally remote compound with biochemical activity with PknB

Follow-up analyses can also take the form of additional prediction rounds that incorporate the results of previous experiments, a setting in which sample-efficiency is paramount. This iterative cycle involving prediction, acquisition, model retraining, and subsequent prediction and acquisition is referred to as “active learning” [28], [31]. In this vein, we conducted a second round of PknB binding affinity predictions after training on both the original dataset and the results from our first round of *in vitro* affinity experiments (**Fig. 3B**). We trained GP and MLP models on this data and again acquired the top five predictions made by each (**Methods**).

All MLP-acquired compounds again had a PknB Kd greater than 10 μM. Although the GP uncertainty scores increased by as much as a factor of 2 from the first round (**Supplementary Fig. 5**), indicating hypotheses that explore riskier, more novel regions of the compound landscape, we still found that one of the GP-acquired compounds, IKK-16, binds PknB with a Kd of 22 nM, the second lowest PknB Kd over all our experiments (**Supplementary Table 2**). IKK-16 was originally developed as an inhibitor of human IκB kinases (IC50 values of 200, 40, and 70 nM for IKKα, IKKβ, and IKK complex, respectively), with *in vivo* activity in an acute murine model of cytokine release [71]. IKK-16 had an acquisition ranking of 24 during the first round but a ranking of 2 in the second round (**Fig. 4D**), indicating that the GP efficiently adapted its beliefs based on a handful of new datapoints to make a successful second-round prediction. Notably, among all training compounds in both the first and second prediction rounds, the most similar structure to IKK-16 is imatinib with a Tanimoto similarity of 0.31 (**Fig. 4E** and **Supplementary Table 4**), indicating that IKK-16 is structurally remote to any compound in the training data; for reference, a recently used novelty threshold was a Tanimoto similarity of 0.40 [12].

We could not find existing literature linking IKK-16 to PknB, suggesting a new potential biomedical dimension for IKK-16 and related small molecules. These results also illustrate how uncertainty combined with an active learning strategy can explore regions of the compound space that are more distal to the original training set and still provide successful predictions.

### Uncertainty prediction improves generative design of novel compound structures

While our discovery experiments leveraged existing, commercially available compounds, our robust predictive models can also help us design new compound structures with high affinity for PknB. In particular, we are interested in a *generative design* paradigm. As in all design problems, a designer wishes to create a new object with a desired property. In machine-assisted generative design, a *generator* algorithm is responsible for generating novel objects while an *evaluator* algorithm prioritizes objects that best fulfill the desired property. In theory, uncertainty prediction enables more robust evaluators that select designs that better reflect the desired, ground-truth property.

We performed generative design of novel small molecule structures that have strong affinity for PknB. Our generation strategy was based on sampling from the latent space of a variational autoencoder (VAE) [72], with an architecture optimized for chemical structures (JTNN-VAE) [43] (**Methods**). We trained a JTNN-VAE to reconstruct the distribution of the entire ZINC/Cayman dataset. We randomly sampled from the JTNN-VAE latent space and decoded the result to obtain an “artificial library” of 200,000 compound structures that do not exist in the ZINC/Cayman dataset (we note that our model for generating chemical structures is distinct from the model used to encode structural features). We used the MLP, MLP + GP, and GP to rank compounds within this artificial library for predicted affinity with PknB, taking uncertainty into account for the latter methods. We emphasize that the generative model learned across the ZINC/Cayman dataset is still highly mismatched from the 72-compound training distribution of the GP, MLP + GP, and MLP. We then used molecular docking, an orthogonal method for binding affinity prediction, to simulate the true binding affinity between the generated compound and the PknB active site. Since consistency across disparate docking scoring functions corresponds to better prediction of true biochemical affinity [73], we use six scoring functions to compare generated designs selected with and without uncertainty [74]–[78].

The molecules prioritized by the GP-based methods had significantly higher affinity than the MLP baseline across all scoring functions, based on one-sided Welch’s *t*-test *P*-values at a false discovery rate (FDR) less than 0.05 [79] (**Fig. 5A**). There was no significant difference between docking scores of GP-based methods and of known high-affinity compounds, used as positive controls (**Fig. 5A**). Visual inspection of the best designs predicted by the GP and by docking reveals structures similar to known inhibitors (**Fig. 5B**), while some of the structures prioritized by the MLP appear pathological (**Fig. 5C**). For the compounds with strong binding affinity, visualizing the binding poses suggested by the docking algorithm shows concordance with a known crystallography-determined small molecule pose [80] (PDB: 2FUM) (**Fig. 5D**). These results show how uncertainty-based robustness in an out-of-distribution setting can better guide the generative design of new chemical structures.

**Figure 5:**
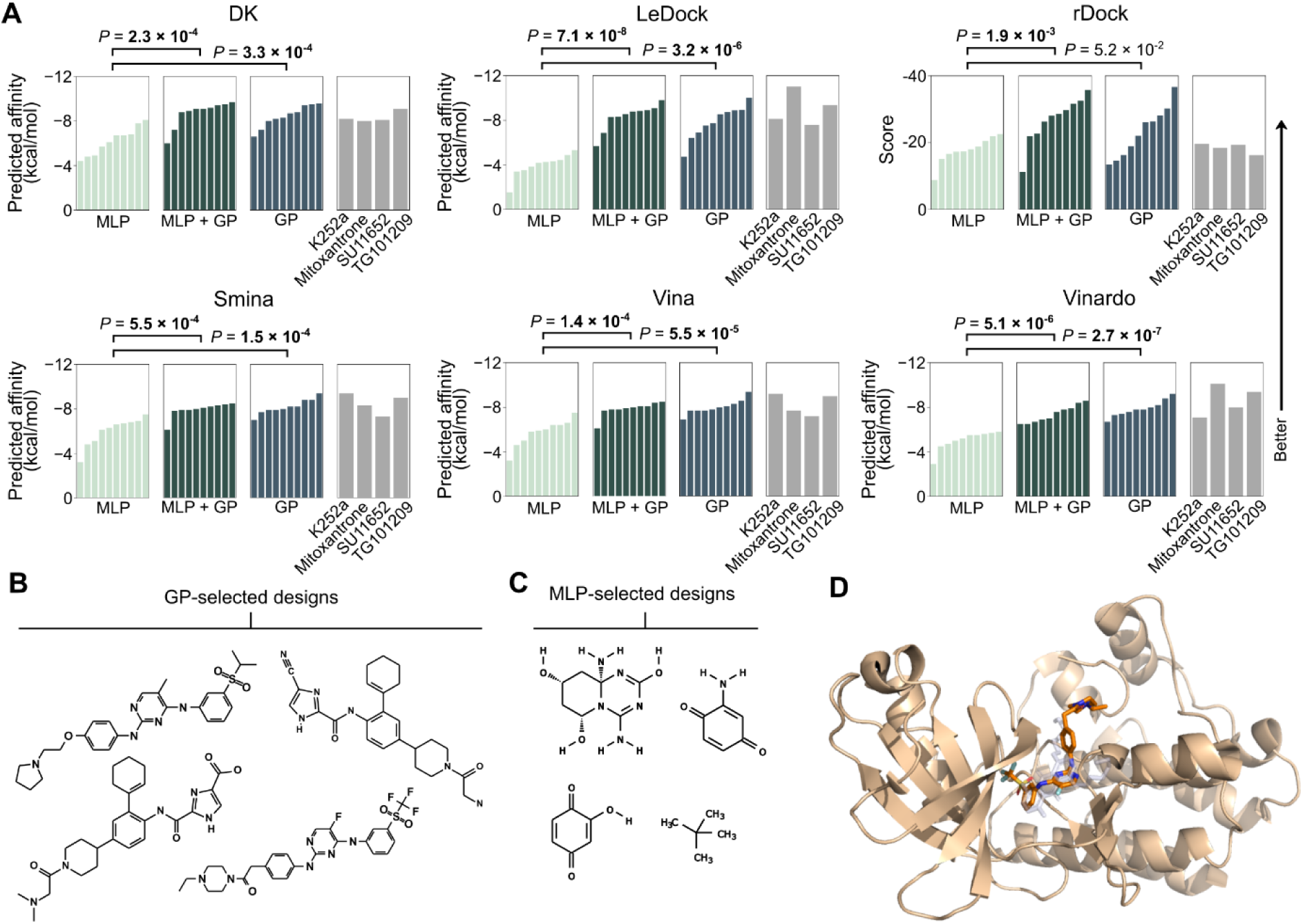
Robust uncertainty prediction for generative design of novel compounds with PknB activity. (**A**) Top ten designs selected by the GP or the MLP + GP have significantly stronger predicted binding affinity compared to the MLP across molecular docking experiments with six different scoring functions. Bolded two-sided independent *t*-test *P*-values indicate statistical significance at FDR < 0.05. Known molecules that bind PknB are provided for comparison; mitoxantrone is the inhibitor with pose determined in the PknB crystal structure in (**D**) (PDB: 2FUM). (**B**) Designs selected by the GP algorithm from a set of 200,000 artificially generated chemical structures resemble known structures that bind PknB (e.g., Fig. 3C). (**C**) Designs selected by the MLP algorithm include pathological structures. (**D**) Visualizing the docking-determined poses of a GP-selected novel design (orange) alongside a mitoxantrone (gray) inhibitor pose determined by X-ray crystallography [80] reveals overlapping locations of certain molecular substructures.

### Generality of uncertainty prediction to disparate biological domains

Importantly, because our approach to machine-guided discovery is general, it can also be applied to diverse domains beyond kinase affinity prediction. Many biological problems, while seemingly disparate, are fundamentally similar in that they are based on predicting the value of target variables based on a set of feature variables (**Fig. 6A**; compare to **Fig. 2B**). We therefore wanted to demonstrate generality by applying our same learning paradigm to predict the brightness of fluorescent proteins based on protein sequence features (**Methods**), potentially enabling an algorithm that can optimize, *in silico*, the fluorescence of an existing protein design [14], [81].

**Figure 6:**
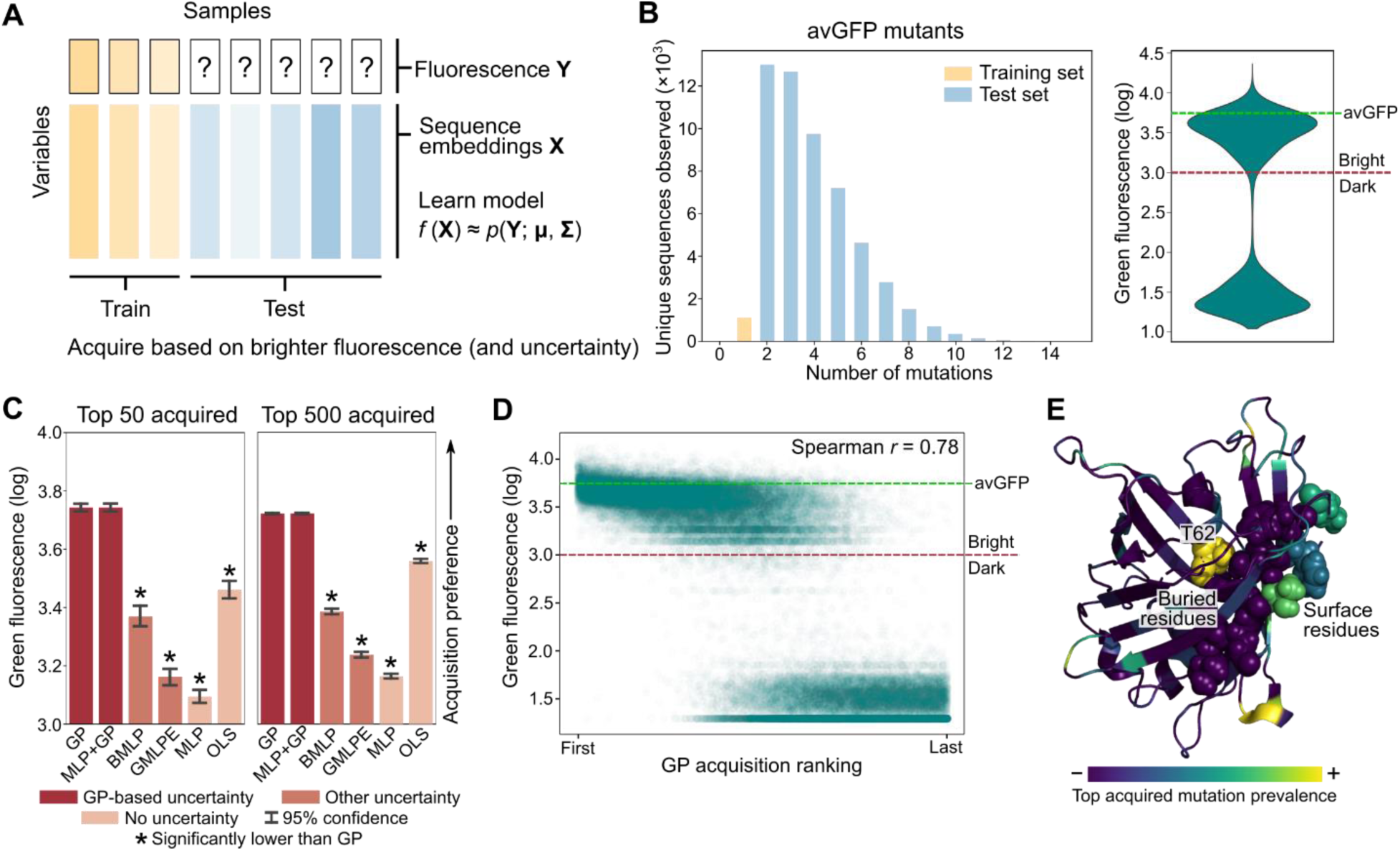
Uncertainty enables robust prediction of protein fluorescence. (**A**) Incorporating uncertainty into predictions is useful in the general setting in which we desire to predict a set of target variables (for example, fluorescence) based on a set of feature variables (for example, protein sequence embeddings); compare to Fig. 2A. (**B**) We train models on avGFP sequences with at most a single-residue mutation compared to wild-type (*n* = 1,115 sequences). We evaluated these models on avGFP sequences with two or more mutations (*n* = 52,910 sequences). The distribution of fluorescence among test set mutants is largely bimodal, with a bright mode and a dark mode; green dashed line indicates median log-fluorescence of wild-type avGFP. (**C**) Models trained on sequences with at most one mutation were used to acquire more highly-mutated sequences, prioritizing higher log-fluorescence (a unitless intensity value). Log-fluorescence was averaged over the top 50 or 500 acquired mutants across five random seeds; bar height indicates mean log-fluorescence. GP-based uncertainty models acquire significantly brighter proteins; statistical significance was assessed with a one-sided *t*-test *P*-value at FDR < 0.05. (**D**) The acquisition ranking produced by the GP is strongly correlated with fluorescent brightness. Each point represents a unique avGFP mutant sequence in the test set. The green dashed line indicates median log-fluorescence of wild-type avGFP. (**E**) In general, GP-acquired mutations are enriched for surface residues over buried residues as expected [2], with T62 being a notable exception. For emphasis, T62 and a beta strand (V93-K101) are displayed as spheres.

We obtained a high-throughput mutagenesis dataset involving avGFP [2]. We trained machine learning models to predict fluorescent brightness based on protein sequence features, where we used the exact same pretrained embedding model as in our kinase experiments [44]. To simulate a data-limited scenario, we only gave the model access to sequences with at most one amino acid mutation compared to wild-type (*n* = 1,115 sequences) (**Fig. 6B**); in contrast, our test set consisted of sequences with two or more mutated residues (*n* = 52,910 sequences), simulating a scenario where an algorithm is asked to make predictions over a more combinatorially complex space. We use the same pretrained learning models as in our kinase cross-validation experiments except for DGraphDTA, a domain-specific model, and CMF, which does not naturally apply to this problem setup; instead, we use ordinary least squares (OLS) regression, a better-suited linear model, as a replacement benchmark (**Methods**).

We once more observed that GP-based models acquired mutant sequences with significantly higher average brightness than other methods (**Fig. 6C**). The GP also performs well in the average case, with an acquisition ranking that is strongly and significantly correlated with measured fluorescence (Spearman *r* = 0.78, two-sided *P* < 10^-308^, *n* = 52,910 sequences) (**Fig. 6D**), which is competitive with or better than baseline methods, many of which overfit the small training dataset (**Supplementary Fig. 6A**). While OLS regression was also robust to overfitting in the average case, GP-based uncertainty results in substantially fewer false positive predictions among top-acquired sequences (**Fig. 6C** and **Supplementary Fig. 6B**). As in the kinase inhibition cross-validation experiments, we observed that GP-based uncertainty helped reduce false-positive predictions among top-acquired samples compared to predictions without uncertainty (**Supplementary Fig. 6C**). These results suggest how predictive (and robust) algorithms could help reduce the complexity of mutagenesis experiments traditionally used to optimize new protein designs [2], [82].

Structurally, many GP-acquired mutations are to surface residues (**Fig. 6E**), consistent with previous observations that buried residue mutations are more likely to be deleterious to fluorescence [2]. A notable exception is high GP prioritization of mutations to T62 (**Fig. 6E**), a buried residue essential to chromophore synthesis [83]. Interestingly, the GP prioritized alanine or serine substitutions (i.e., T62A or T62S) that preserve and even confer higher fluorescence (e.g., a T62A/L178I mutant, the second sequence in the GP acquisition ranking, has a log-fluorescence of 3.9, versus 3.7 for wild-type). While surface residue mutations are more likely to be neutral, an algorithm aimed at enhancement might focus on these buried residues that directly influence fluorescence.

Robust acquisition is useful for highlighting important examples for deeper analysis, and our experiments show that uncertainty improves the robustness of acquisition in different biological settings. We also see that pretrained features and GP-based modeling can improve data-limited performance across disparate biological applications, strengthening the evidence of the broad utility of our learning paradigm.

## Discussion

Biological discovery often requires making educated hypotheses with limited data under substantial uncertainty. In this study, we show how machine learning models that generate biological hypotheses can overcome such challenges. In particular, our results suggest a broadly useful paradigm: neural pretrained features followed by a task-specific supervised GP-based model. We show that uncertainty provides a useful guard against overfitting and pathological model bias, sample efficiency enables successful iterative learning across a broad spectrum of experimental scales, and pretraining elevates our uncertainty models to state-of-the-art performance.

While our study provides concrete illustrations of uncertainty prediction in biochemical activity and protein engineering tasks, uncertainty could also be impactful in many other areas given the complexity of biological structures and systems. The generality of our approach is a notable advantage and stands to benefit from many recent advances in biological representation learning, such as, for example, gene embedding approaches that learn information from coexpression or protein interaction networks [84]. These features can be readily coupled with a GP-based regressor to predict a desirable phenotype; for example, a GP could be trained based on gene embeddings to prioritize CRISPR experiments or other genetic perturbations with the goal of inducing a particular phenotype (for example, cellular fitness or surface marker expression), which is especially useful when acquiring from a combinatorially large interventional search space [85].

Our work highlights GP-based methods as particularly useful. GPs enable theoretically principled incorporation of prior information, can use standard kernels to approximate any continuous function [86], and preserve and even improve modelling performance in complex, nonlinear settings even with limited data. Despite this, GPs have received relatively less attention in learning-based discovery than other methods, like deep learning methods based on large neural network architectures. In our experiments, GP-based methods trained with rich, modern features are sample-efficient and provide well-calibrated uncertainty scores over other nonlinear approaches such as neural ensembles or Bayesian neural networks. A notable methodological finding from our study is that the consistently strong performance of a GP fit to MLP residuals (i.e., MLP + GP) [46] suggests a relatively straightforward way to augment a neural network with uncertainty. There is also much room for methodological development of uncertainty prediction for biological discovery, particularly through related efforts to improve the generalizability of learned models through transfer learning and zero-shot learning [87]–[89].

Importantly, uncertainty-guided prediction provides an efficient alternative, in both time and resources, to large-scale screening or manual trial and error. Focused experimental decision-making is especially important in settings where high-throughput screens are not easy or even tractable. For example, a researcher might first obtain a training dataset with a tractable experiment (for example, a biochemical assay, or a single-gene reporter readout) and follow up a few machine-guided predictions with more complex experiments (for example, involving pathogenic models like Mtb-infected macrophages, or more complex designs like a high-throughput single-cell profiling experiment).

While we mostly focus on “exploitation,” i.e., prioritizing more confident examples that are likelier to yield positive results, researchers can modify the acquisition function to devote more of the experimental budget to “exploration.” With uncertainty, researchers can control the “exploration/exploitation” tradeoff to choose experiments that tolerate a higher risk of failure in order to probe novel regimes [17], [18]. For example, in the drug discovery setting, novelty is important since human-designed drugs are often appraised based on their creativity (in addition to effectiveness) [90], [91]. Novelty is also important across biological domains, such as designing artificial proteins not found in nature [14] or discovering new transcriptional circuits [85]. Uncertainty helps define the boundaries of an algorithm’s knowledge, beyond which human creativity can take over.

Although initializing a model with some training data is helpful, it is also possible to begin with zero training data (all predictions might therefore begin as equally uncertain). As more data is collected, a sample-efficient model with uncertainty can progressively yield better and more confident predictions. This is the iterative cycle of computation and experimentation at the heart of active learning [28], [31], for which we provide a proof-of-concept example in this study.

More generally, we anticipate that iterative experimentation and computation will have a transformative effect on the experimental process. In addition to learning from high-throughput datasets, we also envision learning algorithms working intimately alongside bench scientists as they acquire new data, even on the scale of tens of new datapoints per experimental batch. As we show, using machine learning to generate novel hypotheses will require a principled consideration of uncertainty.

## Acknowledgments

We thank Tristan Bepler and Ellen Zhong for helpful discussions. We thank Diane Ballestas, Patrice Macaluso and Sydney Solomon for assistance with the validation experiments. We thank Robert Chun for assistance with the manuscript. B.H. is partially supported by the Department of Defense (DoD) through the National Defense Science and Engineering Graduate Fellowship (NDSEG) and by NIH grant R01 GM081871 (to B. Berger).

## Declaration of interests

The authors declare no competing interests.

## Methods

### Mycobacterium tuberculosis model details

We utilized wild-type H37Rv and H37Rv expressing an integrated copy of the *luxABCDE* cassette which enables mycobacteria to endogenously produce light [70]; monitoring luminescence of the latter strain has been demonstrated to correlate well with the standard colony forming unit assay [69].

### Human macrophage model details

Human monocytes were isolated from human buffy coats purchased from the Massachusetts General Hospital blood bank using a standard Ficoll gradient (GE Healthcare) and subsequent positive selection of CD14^+^ cells (Stemcell Technologies). Selected monocytes were cultured in ultra-low-adherence flasks (Corning) for 6 days with RPMI media (Invitrogen) supplemented with hydroxyethylpiperazine ethane sulfonic acid (HEPES) (Invitrogen), L-glutamine (Invitrogen), 10% heat-inactivated fetal bovine serum (FBS) (Invitrogen) and 25 ng/mL human macrophage colony-stimulating factor (M-CSF) (Biolegend).

### Multilayer perceptron (MLP)

We trained an MLP with two hidden layers with 200 neurons per layer and rectified linear unit (ReLU) activation functions, trained with mean square error loss and adaptive moment estimation (Adam) with our implementation’s default optimization parameters (learning rate of 0.001, *β*_1_ = 0.9, *β*_2_ = 0.999) [92]. Hyperparameters were tuned based on a small-scale grid search using five-fold random cross-validation within the compound-kinase training set before application to the out-of-distribution test set, with a particular emphasis on preventing overfitting [48]. Lower model capacity, ℓ_2_ regularization of the densely connected layers (weight 0.01), and early stopping after 50 training epochs were helpful in preventing overfitting to the training data and highly pathological outputs on out-of-distribution data (for example, outputting the same prediction value for all instances). The MLP was implemented using the keras Python package (version 2.3.1) using a tensorflow (version 1.15.0) backend with CUDA-based GPU acceleration.

### Collective matrix factorization (CMF)

We performed CMF using the compound-kinase Kds as the explicit data matrix and the neural-encoded compound and kinase features as side-information [53]. Briefly, CMF optimizes the loss function

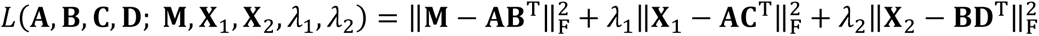

with respect to latent variable matrices **A**, **B**, **C**, and **D**. **M** is the compound-by-kinase binding affinity matrix; **X**_1_ is a side-information matrix where each row contains compound features; **X**_2_ is a side-information matrix where each row contains kinase features; ‖·‖_F_ denotes the Frobenius norm of a matrix; and λ_1_ and λ_2_ are user-specified optimization constants (we set these values to the default value of 1, but observed that cross-validated performance metrics were robust to changes in this parameter). The number of components (i.e., the number of columns in **A**, **B**, **C**, and **D**) was set to the default value of 30, but we also noticed robustness of cross-validated metrics to changes in this parameter. The CMF objective was fit using the limited-memory Broyden–Fletcher–Goldfarb–Shanno algorithm (L-BFGS) via the cmfrec Python package version 0.5.3 [93] (https://cmfrec.readthedocs.io/en/latest/).

### Ordinary least squares (OLS)

For the protein fluorescence experiments, we use OLS as a replacement benchmarking model that is more natural to the problem setup than CMF, which is most commonly used in recommender-system problems involving the affinity between two types of entities (for example, users and shopping items, or compounds and kinases). OLS minimizes the loss *L*(**A**; **X**, **Y**) = 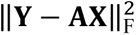 where **Y** is sample-by-target-variable matrix, **X** is the sample-by-feature-variable matrix, and **A** is a learned coefficient matrix (the latter two matrices are also augmented to fit a constant intercept term for each target variable). We use the implementation in the scikit-learn Python package.

### DGraphDTA

We used DGraphDTA [54] to predict compound-kinase Kds. DGraphDTA leverages a graph neural network based on the compound molecular structure and the protein residue contact map. We used the implementation provided at https://github.com/595693085/DGraphDTA with default model architecture hyperparameters. For compound features, we provided the model with chemical SMILE strings that the model transforms into a graph convolutional representation [94]; for kinase features, we use the protein contact maps provided by the original study.

### Prediction acquisition function

For models that output uncertainty scores, an acquisition function is used to rank compound-kinase pairs for acquisition, which in the biological setting often corresponds to further experimental validation, in a way that balances both the prediction value and the associated uncertainty. A standard acquisition function is the upper confidence bound (UCB) [38]. When low prediction values are desirable, UCB acquisition takes the form

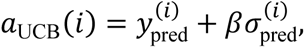

Where 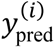 and 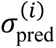 are the predicted Kd and the uncertainty score, respectively, for the *i*th training example and where *β* is a parameter controlling the weight assigned to the uncertainty score. When acquiring the top *k* examples for further experimentation, a researcher can simply take the examples with the *k* lowest values of the acquisition function, i.e., acquire the set

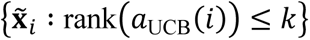

which is a subset of the full unknown test set 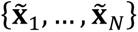. The threshold *k* can reflect an experimental budget (e.g., the size of an experimental batch or the pecuniary cost of an experiment) or it can also be chosen by based on the absolute value of the uncertainty 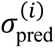 (e.g., only tolerating top-ranked examples with low uncertainty).

### Gaussian Process (GP)

GPs are a Bayesian machine learning strategy that can learn nonlinear functions, can work with limited data, and enable principled incorporation of prior information. The aspect of GPs most relevant to this study is that they enable a researcher to explicitly specify prior information encoding both a “baseline” prediction and corresponding uncertainty. For example, *a priori*, a researcher can assume that a given compound-kinase pair has low affinity; this intuition can be encoded as a probability distribution with most of the probability density assigned to low affinity but with small, nonzero probability assigned to high affinity. On a prediction example that is very different from any training example, the prediction uncertainty of a GP approaches the value of the prior uncertainty [18].

A Gaussian process regressor is fully described by a mean function and a covariance function. For the compound-kinase experiments, our mean function is set to a constant value corresponding to a Kd of 10,000 nM (i.e., the top tested concentration above which the Kd is not determined and a compound-kinase pair is considered inactive). For the protein fluorescence experiments, the mean function is set to a constant value corresponding to a log-fluorescence of 3 (i.e., the original study’s darkness cutoff). Our covariance function is set to a Gaussian, or a squared exponential, kernel scaled by a constant *k*_prior_ related to the prior uncertainty

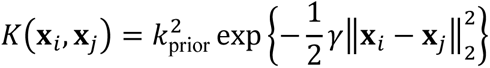

where ‖·‖_2_ denotes the ℓ_2_-distance between feature vectors **x***_i_* and **x***_j_*. For the kinase experiments, *k*_prior_ is set to 10,000 nM; for the protein fluorescence experiments, *k*_prior_ is set to a log-fluorescence of 2; for both, *γ* is set to unity. Each prediction takes the form of a (scalar) Gaussian distribution; we use the mean as the prediction value and the variance as the uncertainty estimate. We use the Gaussian process regressor implementation provided by the scikit-learn Python package. Gaussian processes are reviewed in-depth by Rasmussen and Williams and with helpful, high-level visual aids by Görtler et al.

### Gaussian process fit to residuals of a multilayer perceptron (MLP + GP)

Since much of the interest in machine learning has been on improving the performance of neural network models, a simple way to augment neural networks with uncertainty is to combine the predictions made by a neural network and predictions made by a GP [46]. We use an MLP regressor with the same architecture and hyperparameters as the standalone MLP model described above. The GP fit to the residuals of the MLP regressor has the same form as described for the regular GP above but where the regression problem is formulated as

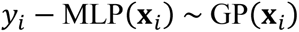

for training example **x***_i_* and training label **y***_i_*. To calculate the prediction value, we evaluate both the MLP and the GP and sum the MLP prediction and the GP mean [46], i.e.,

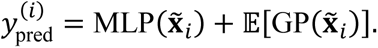

To calculate the uncertainty estimate, we can simply use the GP standard deviation [46], i.e.,

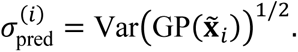

We used the same software (a combination of the scikit-learn, GPyTorch, keras, and tensorflow Python packages) to implement the hybrid model.

### Bayesian multilayer perceptron (BMLP)

A more involved, Bayesian approach to augmenting neural networks with uncertainty is to impose a Bayesian prior on the parameters of the neural network. We train an MLP regressor with the same architecture described above (two hidden layers with 200 neurons per layer and ReLU non linearities) but with a unit-variance Gaussian prior on each weight and bias entry [23]. Within the respective biological task, the Gaussian prior mean for each entry corresponds to a Kd of 10,000 nM (i.e., no biochemical affinity) or a log-fluorescence of 3 (i.e., a dark protein). Optimization was performed under a mean-field independence assumption with gradient descent-based variational inference [23], [47]. When making predictions, we sample 100 neural networks and evaluate each neural network on each prediction example. We use the mean prediction across the 100 neural networks as the prediction value and the variance across the 100 neural networks as the uncertainty estimate. To implement the BMLP, we used the Edward Python package (version 1.3.5) for probabilistic programming [47] with a tensorflow CPU (version 1.5.1) backend.

### Gaussian negative log-likelihood-trained multilayer perceptron ensemble (GMLPE)

Rather than a Bayesian approach to uncertainty, another group of uncertainty methods is based on model ensembles. Ensembling involves fitting multiple models to a training dataset; then, variation in the predictions of the models can be used to estimate uncertainty. For our ensemble method, we use the model described by Lakshminarayanan et al. We train an MLP regressor with the same architecture described above (two hidden layers with 200 neurons per layer and ReLU non linearities) but, instead of mean square error loss, with Gaussian negative log-likelihood loss

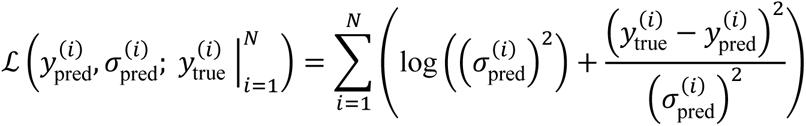

where 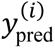 is the predicted value and 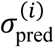 is the predicted uncertainty (both outputted by the neural network), and 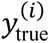 is the ground truth value for training example *i* ∈ {1,2,…*N*}. We train five such models to create a neural network ensemble and we combine prediction distributions across the ensemble as with a Gaussian mixture. As an implementation detail, we trained the neural network to output the log variance to enforce positivity. We implemented the GMLPE with the keras Python package using a tensorflow backend with CUDA-based GPU acceleration.

### Compound-kinase affinity prediction cross-validation setup and benchmarking

We obtained a dataset of binding affinity Kds across all pairs of 72 compounds and 442 kinases (corresponding to 379 unique genes) from Davis et al. Compounds were partitioned randomly into two equal-sized sets of 36 and kinases were partitioned randomly into one set of 190 unique genes (corresponding to 216 kinase proteins, including mutational variants) and another set of 189 unique genes (corresponding to 226 kinase proteins). Compound-kinase pairs therefore fell into one of four “quadrants” of the interaction matrix defined by sets of partitioned compounds and kinases (for a pictorial representation, see **Supplementary Fig. 1A**). One quadrant of the compound-kinase interaction matrix was reserved as training data; the other three quadrants were used as out-of-distribution test data.

Compound structures were obtained from the ZINC database [30] and kinase amino acid sequences were obtained from the UniProt database [95]. For initial computational processing of small molecule structure, we used the RDKit (http://www.rdkit.org/) version 2017.09.1. A compound was featurized based on its structure using a pretrained VAE, optimized for small molecule reconstruction using a graph convolutional junction tree approach (JTNN-VAE), from Jin et al. (https://github.com/wengong-jin/icml18-jtnn). A kinase was featurized based on its amino acid sequence using a pretrained neural language model, designed to encode structural and functional similarities, from Bepler and Berger (https://github.com/tbepler/protein-sequence-embedding-iclr2019). We used the pretrained models made available by both studies without modification. A compound-kinase pair was featurized by the concatenation of the feature vectors of the corresponding compound and kinase. We observed that these state-of-the-art features provided much better empirical performance than traditional features like chemical fingerprints [96] or one-hot-encoded protein family domains [48].

The six benchmarking models (GP, MLP + GP, BMLP, GMLPE, MLP, CMF) were trained on the training quadrant and used to make a prediction (and, if applicable, a corresponding uncertainty score) for each compound-kinase pair in the three test quadrants. Three standard, average-case performance metrics were used: (1) Pearson correlation between predicted and true Kds, (2) Spearman correlation between predicted and true Kds, and (3) mean square error between predicted and true Kds. We also performed a “lead prioritization” experiment. Compound-kinase pairs in the test set were ranked for uncertainty methods by a rank-UCB acquisition function (*β* = 1) and for non-uncertainty methods by the prediction value. We compared the Kds for the top *k* of these compound-kinase pairs across all six methods; we repeated this experiment for *k* = 5 and *k* = 25 to assess different lead prioritization thresholds. The distributions of binding affinities among acquired compound-kinase pairs were assessed for statistical significance (FDR < 0.05) using Welch’s unequal variances *t*-test. The above cross-validation benchmarking experiments (both average-case and lead prioritization) were repeated over five random seeds.

### Acquisition of commercially available ZINC/Cayman compound dataset

We obtained a dataset of small molecule compounds over which we wished to predict binding affinity with various kinases. We used the ZINC database [30], an online repository of chemical compounds with associated metadata for each compound that includes their structure and their commercial availability. We chose the subset of 10,833 compounds in the ZINC database that was also present in the catalog of the Cayman Chemical Company, enabling us to readily purchase high quality compound samples for experimental validation. The only criteria applied to compound selection was that the compound was not present in the Davis et al. training dataset and that the compound was commercially available through the Cayman Chemical Company. We computed statistics over these compounds using the RDKit in Python. ZINC/Cayman compounds were visualized alongside Davis et al. compounds using t-SNE, implemented by the Multicore-TSNE Python package (https://github.com/DmitryUlyanov/Multicore-TSNE). The nearest of neighbor of each ZINC/Cayman compound within the Davis et al. training data was done using Euclidean distance in compound embedding space, using the nearest neighbors implementation in scikit-learn version 0.21.3 [97].

### Experimental validation of compound-kinase affinity predictions

Machine learning models were trained on all compound-kinase pairs from Davis et al. For IRAK4, c-SRC, and p110δ, we trained a GP with high uncertainty weight (*β* = 20) and an MLP. For PknB, the model/acquisition parameter settings were: (1) a GP without considering uncertainty (*β* = 0), (2) a GP with moderate uncertainty weight (*β* = 1), (3) a GP with high uncertainty weight (*β* = 20), (4) an MLP without uncertainty, (5) an MLP + GP with moderate uncertainty weight (*β* = 1), and (6) an MLP + GP with high uncertainty weight (*β* = 20). Compounds from the ZINC/Cayman dataset were featurized using the same pretrained JTNN-VAE as in the cross-validation experiment and concatenated with the feature vector for the corresponding kinase (PknB, IRAK4, c-SRC, or p110δ). Trained models were evaluated on these concatenated features. The top five predictions for each kinase from each of the above models were acquired for binding affinity determination. Predictions involving lipids only commercially available as ethanol solutions were incompatible with the binding assay, excluded from validation, and reported as not interactive.

Compounds were acquired directly from Cayman Chemical. All supplied compounds were tested to ensure ≥ 98% purity. We leveraged the kinase affinity assays provided by the DiscoverX CRO. Kd determination was done using the KdELECT assay, which measures the ability for test compounds to compete with an immobilized, active-site directed ligand using DNA-tagged kinase, where competition is measured via quantitative polymerase chain reaction (qPCR) of the DNA tag. Kinase-tagged T7 phage strains were prepared in an *Escherichia coli* (E. coli) host derived from the BL21 strain. E. coli were grown to log-phase and infected with T7 phage and incubated with shaking at 32°C until lysis. The lysates were centrifuged and filtered to remove cell debris. Streptavidin-coated magnetic beads were treated with biotinylated ligand for 30 minutes at room temperature to generate affinity resins for kinase assays. The liganded beads were blocked with excess biotin and washed with blocking buffer [SeaBlock (Pierce), 1% bovine serum albumin (BSA), 0.05% Tween 20, 1 mM dithiothreitol (DTT)] to remove unbound ligand and to reduce non-specific binding.

Binding reactions were assembled by combining kinases, liganded affinity beads, and test compounds in 1X binding buffer [20% SeaBlock, 0.17X phosphate-buffered saline (PBS), 0.05% Tween 20, 6 mM DTT]. Test compounds were prepared as 111X stocks in 100% DMSO. Kds were determined using an 11-point 3-fold compound dilution series with three DMSO control points with a top test compound concentration of 10,000 nM. All compounds for Kd measurements are distributed by acoustic transfer (non-contact dispensing) in 100% DMSO. The compounds were then diluted directly into the assays such that the final concentration of DMSO was 0.9%. All reactions performed in polypropylene 384-well plate. Each was a final volume of 0.02 mL. The assay plates were incubated at room temperature with shaking for 1 hour and the affinity beads were washed with wash buffer (1x PBS, 0.05% Tween 20). The beads were then re-suspended in elution buffer (1x PBS, 0.05% Tween 20, 0.5 μM non-biotinylated affinity ligand) and incubated at room temperature with shaking for 30 minutes. The kinase concentration in the eluates was measured by qPCR.

Kds were calculated with a standard dose-response curve using the Hill equation

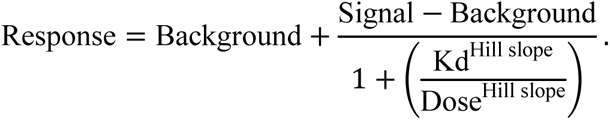

Curves were fitted using a non-linear least square fit with the Levenberg-Marquardt algorithm. The Hill slope was set to -1; a deviation from this Hill slope in the dose-response pattern was used to identify possible aggregation, but no such deviation was observed. A full dose-response curve was obtained and fit in duplicate, and we report the mean Kd between duplicate curves.

### Axenic Mtb growth inhibition assay

H37Rv Mtb growth was evaluated using the resazurin viability assay (alamar blue). Mtb was grown to an optical density (OD) corresponding to early log phase (OD 0.4) and back-diluted to an optical density of 0.003 in 7H9 media supplemented with oleic albumin dextrose catalase (OADC) prior to incubation with a range of concentrations of K252a, TG101209, SU11652, and rifampicin or vehicle control in a 96 well plate with shaking at 37°C. Bacteria were incubated with drug alone for 72 hours prior to the addition of alamar blue. After addition of alamar blue, H37Rv was incubated for an additional 48 hours and alamar blue absorbance was measured using a Tecan Spark 10M. Normalized alamar blue absorbance was calculated as

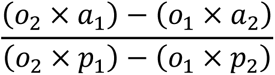

where *o*_1_ = 80586 is the molar extinction coefficient of oxidized alamar blue at 570 nm; o_2_ = 117216 is the molar extinction coefficient of oxidized alamar blue at 600 nm; *a*_1_ and *a*_2_ are the measured absorbance of the test well at 570 nm and 600 nm, respectively; and *p*_1_ and *p*_2_ are the measured absorbance of a positive growth control well at 570 nm and 600 nm, respectively. For each compound, we assessed bacterial growth at 1.25, 2.5, 5, 10, 25, and 50 μM to determine the MIC.

Additionally, Mycobacterium tuberculosis strain H37Rv bacteria expressing an integrated copy of the *luxABCDE* cassette [70] were grown to mid-log phase and diluted to an optical density of 0.006. Mycobacteria were added to wells of a 96-well solid white polystyrene plate and incubated with a vehicle control (DMSO) or rifampicin, TG101209, or SU11652 (Cayman Chem) for 5 days. Plates were sealed with breathable film (VWR) and incubated at 37°C for 4 days with shaking. On day 5, we measured luminescence as a proxy for total bacterial burden.

### Primary Human Macrophage Culture

Deidentified buffy coats from healthy human donors were obtained from Massachusetts General Hospital. Peripheral blood mononuclear cells (PBMCs) were isolated from buffy coats by density-based centrifugation using Ficoll (GE Healthcare). CD14^+^ monocytes were isolated from PBMCs using a CD14 positive-selection kit (Stemcell). Isolated monocytes were differentiated to macrophages in RPMI 1640 (ThermoFisher Scientific) supplemented with 10% heat-inactivated fetal FBS (ThermoFisher Scientific), 1% HEPES, and 1% L-glutamine. Media was further supplemented with 25 ng/mL M-CSF (Biolegend, MCSF: 572902). Monocytes were cultured on low-adhesion tissue culture plates (Corning) for 6 days. After 6 days, macrophages were detached using a detachment buffer of 1X Ca-free PBS and 2 mM ethylenediaminetetraacetic acid (EDTA), pelleted, and recounted. Macrophages were plated in tissue culture-treated 96-well solid white polystyrene plates at a density of 50,000 cells per well in maintenance media (RPMI, 10% heat-inactivated FBS, 1% HEPES, and 1% L-glutamine) and allowed to re-adhere overnight.

### Intra-macrophage Mtb growth inhibition assay

H37Rv Mtb expressing the *luxABCDE* cassette were grown to an optical density of 0.4 and centrifuged briefly. Mtb were resuspended in pre-warmed maintenance media and filtered through a 5 μM filter to remove clumped bacteria and generate a single-cell suspension. Macrophages were infected at a multiplicity of infection of 3 bacteria to 1 macrophage in 100 μL per well and phagocytosis was allowed to proceed for 4 hours prior to washing macrophages twice with pre-warmed maintenance media to remove extracellular bacteria. Following phagocytosis and washing, cells were incubated with media containing a DMSO vehicle control or rifampicin, K252a, or SU11652 (Cayman Chem) for 5 days. On day 5, we measured luminescence as a proxy of intracellular bacterial burden as previously described [69], [70] using a high-throughput luminometer.

### Second Round of Acquisition and Validation of Compound-PknB Interactions

A GP or an MLP was trained on both the original training dataset [1] and all of the first-round PknB-related experimental data (**Fig. 3B**). All other training and acquisition details were the same as in the first prediction round. The top five predictions for the GP (*β* = 20) and for the MLP (ten predictions in total) had their binding affinity Kds determined via the same *in vitro* binding assay described above. IKK-16, acquired by the GP, was the sole hit with Kd below 10,000 nM. To assess the similarity between IKK-16 and each of the original training set compounds, we used the RDKit to compute the Tanimoto similarity of Morgan fingerprints with a radius of 2 and 2048 bits, the same Tanimoto similarity computation procedure as in Stokes et al.

### Generation of artificial compound library and affinity prediction

We used a machine-learning method to generate a library of 200,000 unique compound structures not present in the ZINC/Cayman database. To do so, we trained a JTNN-VAE [43] to reconstruct the ZINC/Cayman dataset of 10,833 compounds using the default model architecture parameters in the publicly available training code (https://github.com/wengong-jin/icml18-jtnn). Hyperparameters were selected based on the provided defaults, namely an embedding dimension of 56, a batch size of 40, a hidden dimension of 350, a depth of 3, a learning rate of 0.001, and termination of training after 40 epochs. To generate new compounds, random vectors were sampled uniformly across the 56-dimensional latent space and then decoded by the JTNN-VAE; molecule structures present in the training data were discarded and not considered as novel designs. We note that the JTNN-VAE model for chemical featurization is a different, pretrained model used in the original study [43]; we trained a separate JTNN-VAE model for molecule generation. These artificially generated compound structures were featurized and concatenated with the PknB feature vectors as described previously. We then acquired the top ten compounds according to a GP (*β* = 20), an MLP + GP (*β* = 20), and an MLP, where all models were trained exclusively on the Davis et al. kinase inhibition data, as described previously.

### Docking-based validation of compound designs

We used a crystallography-determined structure of PknB in complex with mitoxantrone (PDB: 2FUM) [80] as the underlying structure for our docking procedure. To prepare the kinase structure for docking, we restricted our analysis to chain A with the mitoxantrone molecule removed. Our docking region encompassed the full set of amino acids directly proximal to the ligand pocket (https://www.rcsb.org/3d-view/2FUM?preset=ligandInteraction&sele=MIX). Structure files were preprocessed to be in compatible file formats using AutoDockTools version 1.5.6 and Open Babel version 2.3.2 [98]. We used LeDock version 1.0 [76], rDock version 2013.1 [77], AutoDock Vina version 1.1.2 [78], and smina version “Oct 15 2019” [75], the last of which implemented the DK and Vinardo [74] scoring functions in addition to its own default scoring function. The Vina- and smina-based toolkits were run with an exhaustiveness parameter of 500, a high value meant to increase the search space of possible poses, and all other parameters set to the default. We use the reported energy scores returned by the docking procedure. We visualized docking poses using PyMOL version 2.3.3.

### Protein fluorescence cross-validation setup and benchmarking

We obtained mutagenesis data from Sarkisyan et al., treating each unique avGFP sequence as a separate sample featurized by embeddings derived from the full protein sequence. We used the same pretrained sequence embedding model from Bepler and Berger [44] used to featurize kinase sequences in our other experiments. The original mutagenesis study assigned a median log-fluorescence value to each unique sequence, obtained via fluorescence-activated cell sorting of GFP-expressing bacterial vectors based on brightness at 510 nm emission [2]. Our supervised formulation is to predict log-fluorescence based on the neural sequence embedding. We use a log-fluorescence of 3 as a cutoff (as used in the original study) below which all sequences are considered equally dark (i.e., when training the model, we set all dark sequences to a log-fluorescence value of 3).

GP, MLP + GP, BMLP, GMLPE, MLP, and OLS regressors were each trained on 1,115 unique sequences containing at most one mutation to wild-type avGFP (UniProt: P42212). After training, we acquired sequences among the remaining avGFP mutants (a total of 52,910 unique sequences) from the same study based on higher predicted brightness, and, if available, low predicted uncertainty using rank-UCB (*β* = 1). We compared models based on the top 50 or 500 acquired sequences; models with pseudo-randomness were run across five random seeds. The distributions of fluorescence among acquired sequences were assessed for statistical significance (FDR < 0.05) using Welch’s unequal variances *t*-test. We also measured the Spearman correlation between acquisition rank and each mutant sequence’s median log-fluorescence.

Mutations involved in the top hundred acquired sequences by the GP were located on an X-ray crystallography-determined avGFP structure (PDB: 2WUR) [99]. We used the FindSurfaceResidues (https://pymolwiki.org/index.php/FindSurfaceResidues) PyMOL script to distinguish buried and surface residues. We used PyMOL to visualize the protein structure.

### Benchmarking hardware and computational resources

Experiments had access to a Nvidia Tesla V100 PCIe GPU (32GB RAM) and an Intel Xeon Gold 6130 CPU (2.10GHz, 768GB of RAM). GP training for kinase and GFP experiments required approximately 60 and 30 minutes of runtime, respectively. All experiments required a maximum of 50 GB of CPU RAM or 32 GB of GPU RAM.

### Statistical analysis implementation

We use the scientific Python toolkit, including the scipy (version 1.3.1) and numpy (version 1.17.2) Python packages [100], to compute the statistical tests described in the manuscript, including Pearson correlation, Spearman correlation, Welch’s unequal variance *t*-test, and associated *P* values. We use the seaborn Python package (version 0.9.0; https://seaborn.pydata.org/) to compute the 95% confidence intervals and violin-plot kernel density estimates in our data visualizations.

### Data and code availability

We make our data and code available at http://cb.csail.mit.edu/cb/uncertainty-ml-mtb/.

## Notes

### Competing Interest Statement

The authors have declared no competing interest.

