## Supplementary Information for "Learning with uncertainty for biological discovery and design"

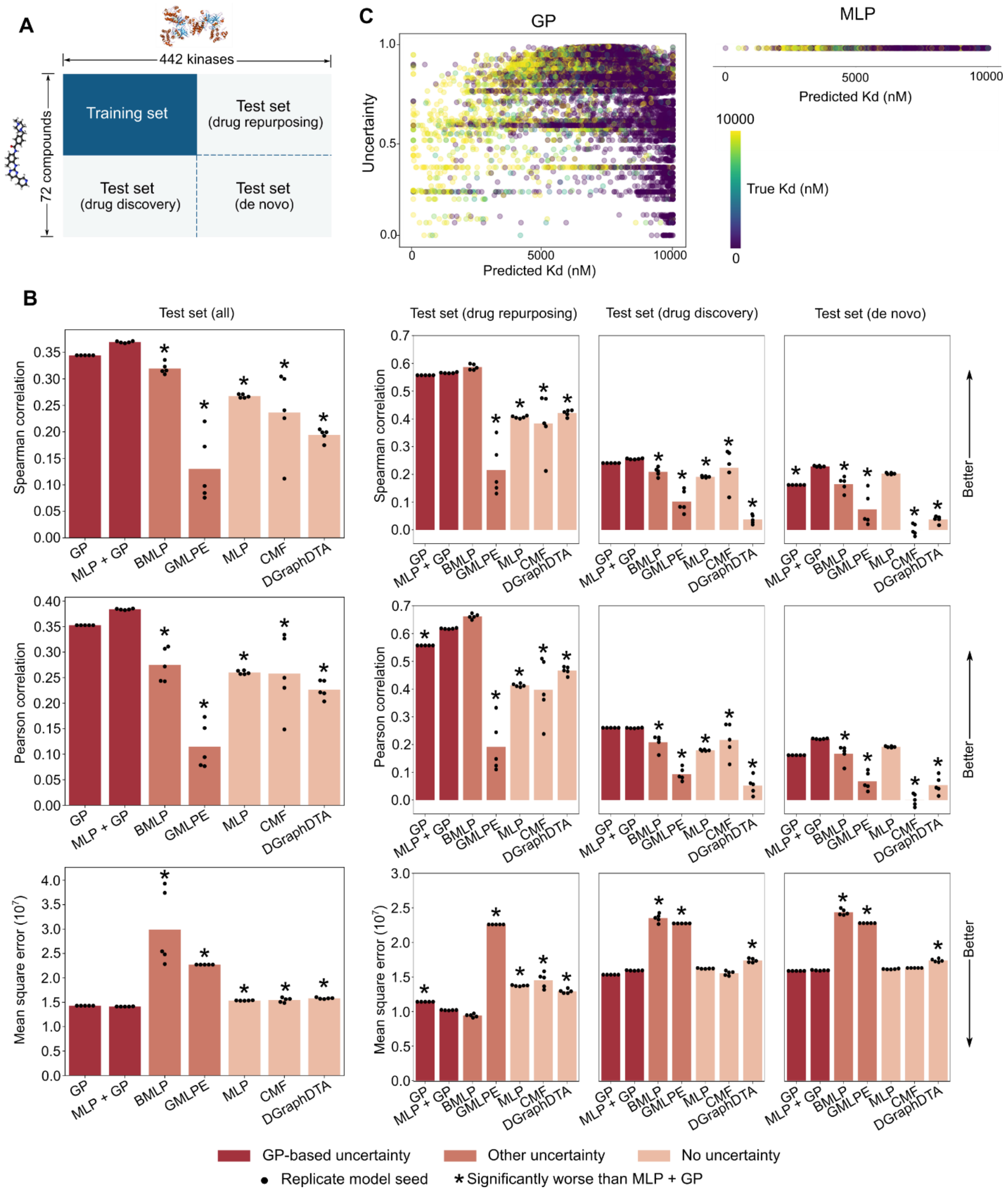


**Supplementary Figure 1: Performance of machine learning methods on out-of-distribution cross validation experiments, related to Figure 2**

(**A**) For cross validation experiments, we used a dataset of compound-kinase Kds among all pairs of 72 compounds and 442 kinases from Davis et al. The dataset was partitioned such that portions of the test set would have compounds not seen in the training data ("drug discovery"), kinases not seen in the training data ("drug repurposing"), and entire compound-kinase pairs not seen in the training data ("de novo"). (**B**) Performance within each out-of-distribution test set quadrant (**A**) was measured with average-case metrics (Spearman correlation, Pearson correlation, or mean square error between model predictions and ground truth Kds). Bar height indicates mean. (**C**) Scatter plots show the relationship between predicted Kd and model uncertainty, colored by the ground truth Kd, for all items in the test set. GP uncertainty scores (scaled to be between 0 and 1, inclusive) that are lower also correspond to better separation of active and inactive interactions. A normal MLP does not output uncertainty estimates. Without quantifying uncertainty, predictions may be overconfident.

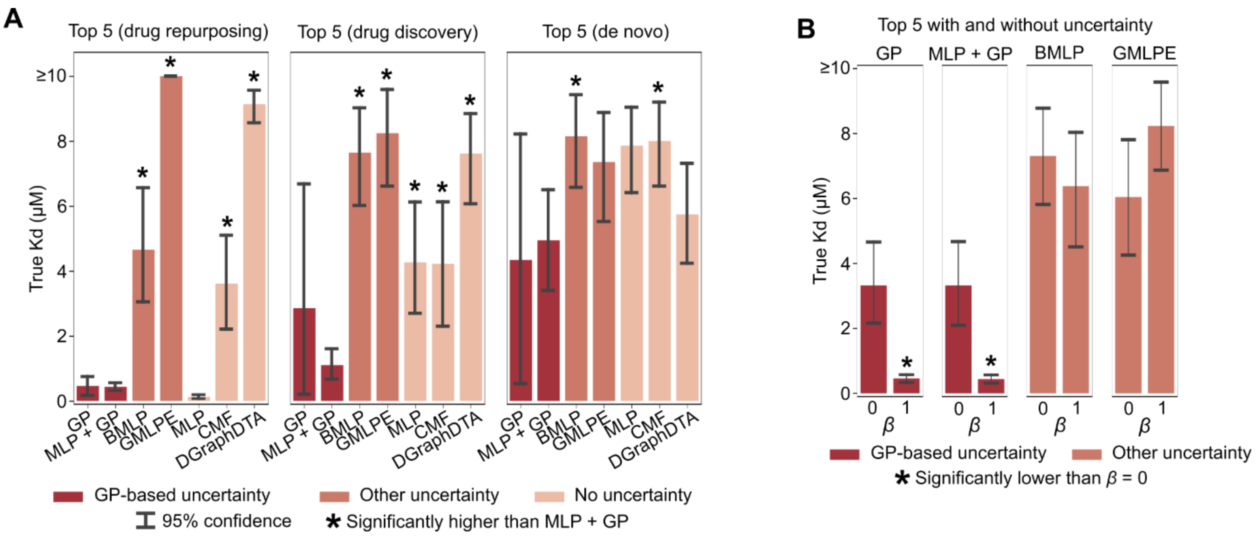


**Supplementary Figure 2: Lead prioritization statistics from cross validation experiments, related to Figure 2**

(**A**) Performance within each out-of-distribution test set quadrant was measured based on lead-prioritization (the true Kd of the top 5 acquired compounds in each random seed). Bar height indicates mean; statistical significance was assessed with a one-sided Welch's *t*-test *P*-value (FDR < 0.05). (**B**) The true Kd of the top 5 acquired compounds for each uncertainty model with ($\beta = 1$) and without uncertainty ($\beta = 0$). Bar height indicates mean; statistical significance was assessed with a one-sided Welch's *t*-test *P*-value (FDR < 0.05).

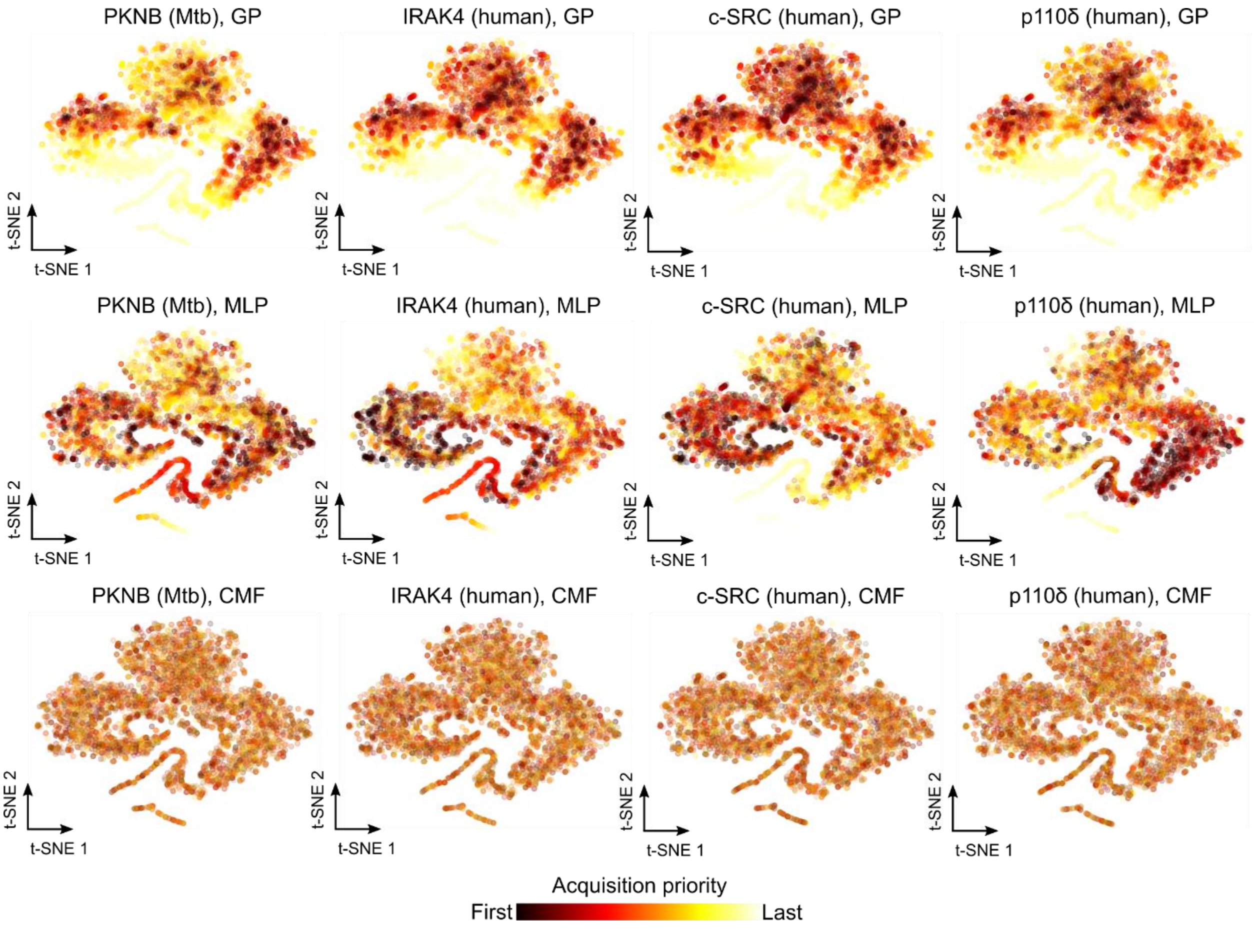


**Supplementary Figure 3: Visualization of ZINC/Cayman acquisition priority, related to Figure 2**

Each compound in the ZINC/Cayman library is visualized as a two-dimensional t-SNE of the chemical embedding space, colored according to acquisition priority for high predicted binding affinity (and, if available, low uncertainty) to four kinases. GP-based acquisition prioritizes regions of the compound space close to available training data (**Figures 2F** and **2G**). In contrast, MLP-based acquisition consistently prioritizes compounds that are out-of-distribution, indicating potentially pathological predictions. CMF predictions appear to lack any meaningful structure with regards to the compound landscape. PknB visualizations are the same as in **Figure 2** and reproduced here for comparison.

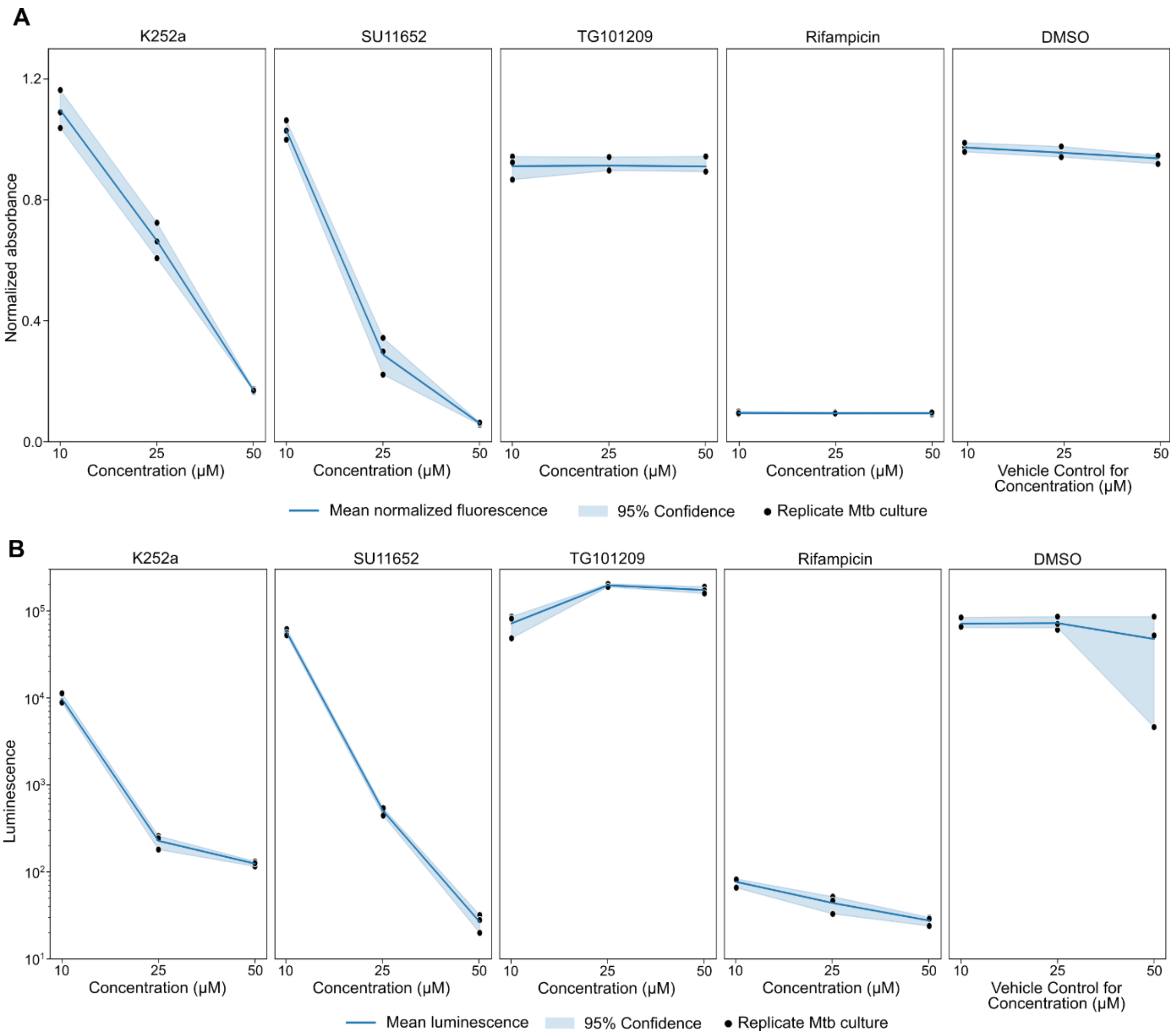


**Supplementary Figure 4: Axenic Mtb growth dose-response, related to Figure 4**

(**A**) Growth of axenic Mtb measured via alamar blue absorbance normalized to untreated condition after five days of incubation in the presence of PknB-inhibiting compounds (K252a, SU11652, and TG101209), a known antibiotic (Rifampicin), and a vehicle control (DMSO). (**B**) Growth of axenic luciferase-expressing Mtb measured as luminescence after five days of incubation in the presence of PknB-inhibiting compounds (K252a, SU11652, and TG101209), a known antibiotic (Rifampicin), and a vehicle control (DMSO).

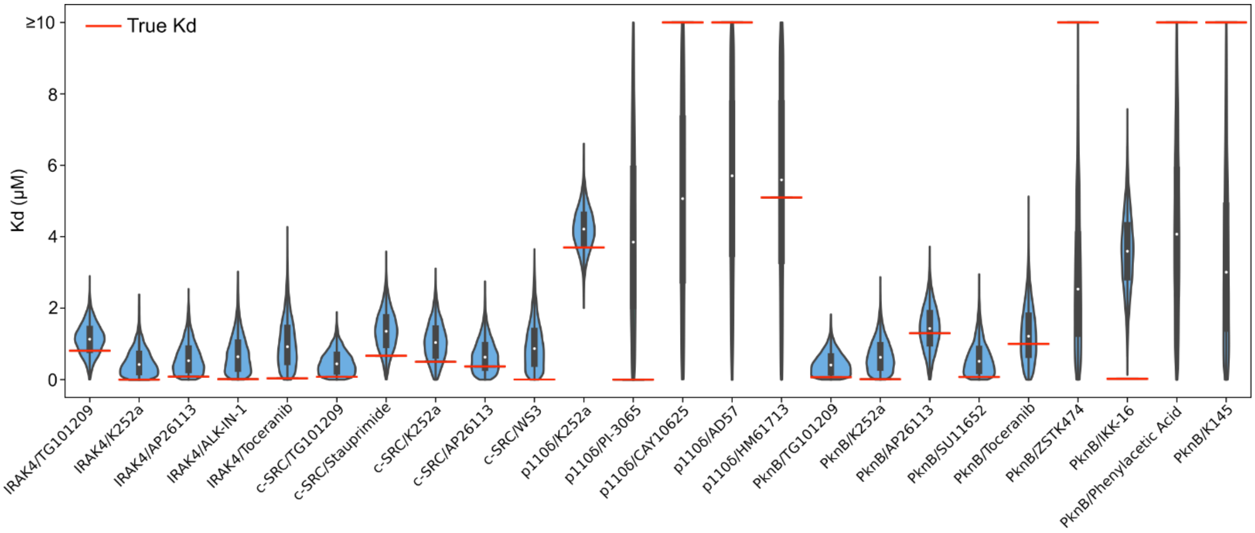


**Supplementary Figure 5: Prediction uncertainty distributions and true values, related to Figures 3 and 4**

Violin plots and box plots correspond to the GP output for a given compound/kinase pair; the box extends from the first to third quartile, the whiskers extend from the min to max, and the white dot indicates the median. Horizontal red lines correspond to the true experimentally determined Kd. Note that uncertainty in addition to the prediction value adds interpretability; for example, the GP-outputted distributions corresponding to p110δ/K252a and PknB/Phenylacetic acid have similar means but different variances, with greater tolerance for a false positive prediction in the latter.

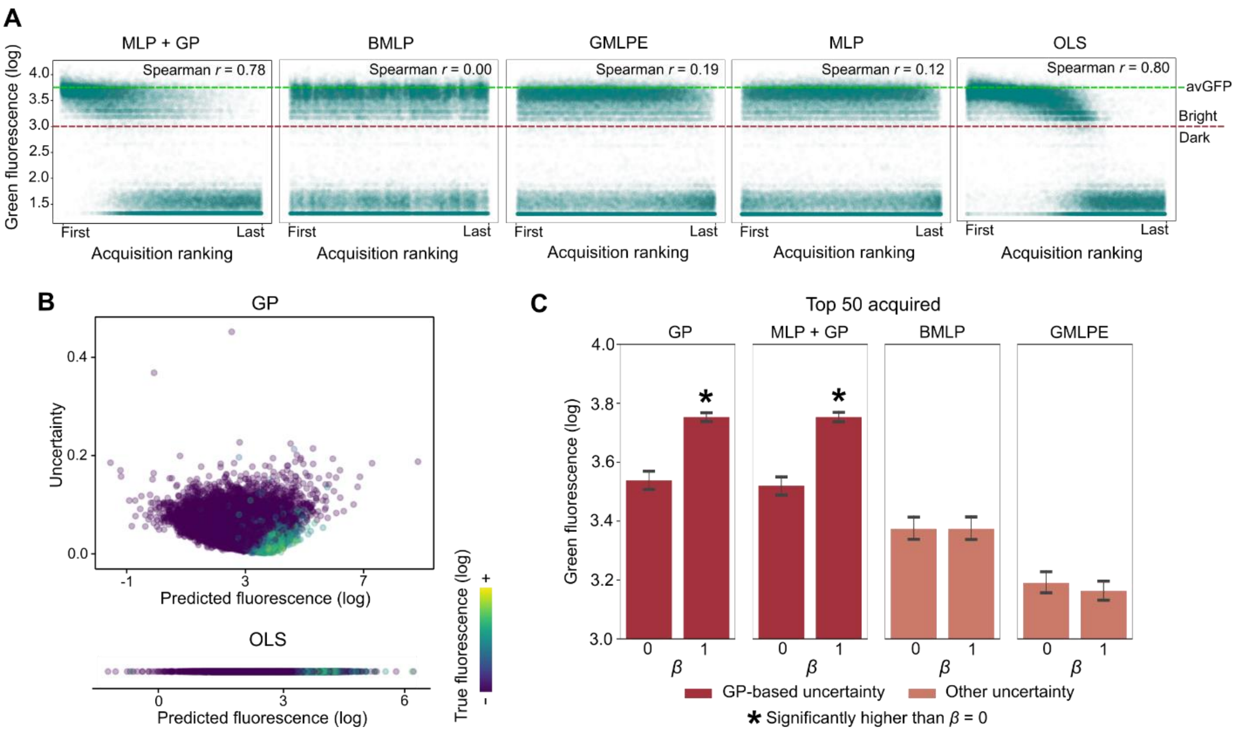


**Supplementary Figure 6: Comparison of model acquisition of avGFP mutants, related to Figure 6**

(**A**) Scatter plots showing true log-fluorescence versus acquisition ranking for benchmarked machine learning methods; compare to the same plot for our GP model in **Figure 6D**. Neural network-based models (without GP augmentation) have low correlation between acquisition ranking and fluorescence, perhaps due to overfitting. OLS has comparable correlation to GP, but significantly worse performance compared to GP-based models among the top ranked examples (**Figure 6C**). Each point corresponds to a unique avGFP mutant; the green dashed line indicates median log-fluorescence of wild-type avGFP. (**B**) Each point corresponds to a test set mutant sequence. At lower uncertainty scores, the GP separates truly bright fluorescing sequences from truly dark fluorescing sequences; a similar pattern was observed for the GP + MLP. OLS regression does not compute uncertainty scores, leading to overconfident predictions among top-acquired sequences. (**C**) The true log-fluorescence of the top 50 acquired mutant sequences for each uncertainty model with ($\beta = 1$) and without uncertainty ($\beta = 0$). Bar height indicates mean; statistical significance was assessed with a one-sided Welch's *t*-test *P*-value (FDR < 0.05).

| Property | Number of Samples | Minimum | Median | Maximum | Mean | Standard Deviation |
|---|---|---|---|---|---|---|
| Exact molecular weight (Da) | 10,833 | 61.0 | 352.2 | 994.5 | 367.9 | 140.3 |
| SSSR | 10,833 | 0 | 2 | 12 | 2.4 | 1.7 |
| Balaban J | 10,833 | 0.7 | 2.0 | 6.3 | 2.2 | 0.8 |
| Bertz CT | 10,833 | 17.2 | 661.3 | 2850.1 | 734.0 | 399.1 |
| Tanimoto similarity (RDK Fingerprint, 2048 bits) | 58,671,528 | 0.00 | 0.18 | 1.00 | 0.20 | 0.11 |
| Tanimoto similarity (Morgan Fingerprint, 2048 bits, radius = 2) | 58,671,528 | 0.00 | 0.09 | 1.00 | 0.11 | 0.07 |

**Supplementary Table 1: Statistics for ZINC/Cayman dataset, related to Figure 2.**

Various statistics computed over the 10,833 chemicals in the ZINC/Cayman dataset, namely, exact molecular weight in Daltons (Da), the size of the smallest set of smallest rings (SSSR), and measurements of molecular complexity (Balaban's J value and Bertz's CT value). Statistics of chemical similarity, namely Tanimoto similarity of fingerprints produced by two different fingerprint methods, were computed over all 58,671,528 combinations of chemicals.

| **Compound** | **PknB** | **IRAK4** | **c-SRC** | **p110δ** |
|---|---|---|---|---|
| (1'S,2'S)-Nicotine-1'-oxide | >10000 | | | |
| 3-O-methyl-N-acetyl-D-Glucosamine | | | >10000 | |
| 8-iso Prostaglandin $E_2$ | | | >10000 | |
| AB-BICA | >10000 | | | |
| Abacavir | | >10000 | | |
| AD57 | | | | >10000 |
| ALK-IN-1 | 620 | 13 | | |
| AP26113 | 1300 | 83 | 370 | |
| Atracurium | >10000 | | | |
| AVL-292 | 5500 | | | |
| CAY10625 | | | | >10000 |
| Epinastine | >10000 | | | |
| Evodiamine | | | >10000 | |
| GLYX 13 | | | >10000 | |
| GP-NEPEA | | | | >10000 |
| HM61713 | | | | 5100 |
| IKK-16 | 22 | | | |
| K145 | >10000 | | | |
| K252a | 11 | 0.85 | 500 | 3700 |
| Lovastatin | >10000 | | | |
| LY2886721 | >10000 | | | |
| Mevastatin | >10000 | | | |
| NVP-TAE226 | 9900 | | | |
| Oxymatrine | >10000 | | | |
| Phenylacetic Acid | >10000 | | | |
| PI-3065 | | | | 0.36 |
| PX 1 | >10000 | | | |
| Ro 4929097 | >10000 | | | |
| Ro 67-7476 | | | >10000 | |
| S-(5'-Adenosyl)-L-methionine chloride | >10000 | | | |
| Stauprimide | | | 670 | |
| SU11652 | 76 | | | |
| TG101209 | 71 | 810 | 79 | |
| Toceranib | 1000 | 37 | | |
| WAY-161503 | | >10000 | | |
| WS3 | | | 4 | |
| ZSTK474 | >10000 | | | |

**Supplementary Table 2: Summary of tested interactions, related to Figures 3 and 4**

All Kds determined in this study in units of nM. Blank cells indicate interactions that were not acquired. “>10000” indicates a Kd greater than the top concentration of 10,000 nM. Cells with a Kd of 100 nM or less are highlighted in blue.

| Compound | MIC (µM) |
|---|---|
| K252a | 25 |
| Rifampicin | 1.25 |
| SU11652 | 25 |
| TG101209 | >50 |

**Supplementary Table 3: MIC values from axenic culture experiments**

MIC values determined by assessing bacterial growth in the presence of compounds at 1.25, 2.5, 5, 10, 25, and 50 µM via an alamar blue assay.

| Acquired compound | Target (Kd < 100 nM) | Closest compound | Morgan fingerprint Tanimoto similarity |
|---|---|---|---|
| IKK-16 | PknB | Imatinib | 0.31 |
| PI-3065 | p110δ | GDC-0941 | 0.46 |
| ALK-IN-1 | IRAK4 | TAE-684 | 0.55 |
| TG101209 | PknB, c-SRC | TG-101348 | 0.68 |
| Toceranib | PknB, c-SRC | Sunitinib | 0.72 |
| AP26113 | IRAK4 | TAE-684 | 0.73 |
| WS3 | c-SRC | AST-487 | 0.73 |
| K252a | PknB, IRAK4 | CEP-701 | 0.77 |
| SU11652 | PknB | Sunitinib | 0.81 |

**Supplementary Table 4: Closest original training set compounds, related to Figure 4**

For each acquired compound involved in a potent binding interaction (Kd < 100 nM; see **Table S2**), the third column from the left reports the most similar compound in the training data and the fourth column reports the Tanimoto similarity.
